# A genome-wide association screen for genes affecting leaf trichome development and epidermal metal accumulation in Arabidopsis

**DOI:** 10.1101/2024.09.10.612273

**Authors:** Radek Bezvoda, Yazmín Mónica Landeo-Ríos, Zdeňka Kubátová, Eva Kollárová, Ivan Kulich, Wolfgang Busch, Viktor Žárský, Fatima Cvrčková

**Affiliations:** Department of Experimental Plant Biology, Faculty of Sciences, Charles University, Prague, Czechia; Plant Molecular and Cellular Biology Laboratory, and Integrative Biology Laboratory, Salk Institute for Biological Studies, La Jolla, CA, USA; Gregor Mendel Institute (GMI), Austrian Academy of Sciences, Vienna Biocenter (VBC), Vienna, Austria; Institute of Experimental Botany, Czech Academy of Sciences, Prague, Czechia

**Keywords:** *Arabidopsis thaliana*, GWAS, phenotypic variability, trichome, guard cell, metal accumulation, BioClim

## Abstract

To identify novel genes engaged in plant epidermal development, we characterized the phenotypic variability of rosette leaf epidermis of 310 sequenced *Arabidopsis thaliana* accessions, focusing on trichome shape and distribution, compositional characteristics of the trichome cell wall, and histologically detectable metal ion distribution. Some of these traits correlated with climate parameters of ourq accession’s locations of origin, suggesting environmental selection. A novel metal deposition pattern in stomatal guard cells was observed in some accessions. Subsequent GWAS analysis identified 1546 loci with protein sequence-altering SNPs associated with one or more traits, including 5 genes with previously reported relevant mutant phenotypes and 80 additional genes with known or predicted roles in relevant developmental and cellular processes. Some candidates, including GFS9/TT9, exhibited environmentally correlated allele distribution. Several large gene families, namely DUF674, DUF784, DUF1262, DUF1985, DUF3741, cytochrome P450, receptor-like kinases, Cys/His-rich C1 domain proteins and formins were overrepresented among the candidates for various traits, suggesting epidermal development-related functions. A possible participation of formins in guard cell metal deposition was supported by observations in available loss of function mutants. Screening of candidate gene lists against the STRING interactome database uncovered several predominantly nuclear protein interaction networks with possible novel roles in epidermal development.

## Introduction

The plant above-ground organs epidermis is a sophisticated interface between the plant and its environment, providing mechanical protection, deterring pathogens and predators and allowing for a controlled gas exchange via stomata. Multiple cell types act in a concerted manner to fulfil these functions, and even subtle disruption of their differentiation may cause observable phenotypic alterations (Zuch et al., 2022). Unconspicuous epidermal defects, well tolerated under laboratory conditions, can indicate perturbation of fundamental cell differentiation or cell morphogenesis processes, as documented, for example, by identification of mutants defective in actin nucleation based on a distorted trichome phenotype (see Szymanski, 2005).

The most abundant cell type in the leaf epidermis of the model plant *Arabidopsis thaliana* are the puzzle piece-shaped pavement cells. Their morphogenesis is driven by microtubule rearrangements that constrict cell surface expansion to generate indentations and allow actin-dependent intrusion of neighboring cell’s lobes. Coordination of these processes involves cell wall-mediated mechanical feedback and is orchestrated by a regulatory network including ROP small GTPases, their effectors, localized exo- and endocytosis and auxin signaling (see Sapala et al., 2018; Lin and Yang, 2020; Liu et al., 2021; Igisch et al., 2022). Leaf epidermis also contains stomatal complexes, hydathodes and trichomes, whose patterning is controlled by complex regulatory networks (see Torii, 2021). Differentiation of these specialized cell types involves rearrangement of gene expression patterns, well documented in *A. thaliana* for both stomatal guard cells (Zhao et al., 2008; Bates et al., 2012; Xia et al., 2022) and trichomes (Jakoby et al., 2008; Huebbers et al., 2022, Huebbers et al., 2023).

Trichomes contribute to anti-herbivore defense by acting as mechanical deterrents of herbivore feeding; their ecological relevance is well documented (e.g. Sato et al., 2019; Qu et al., 2022). They also engage in detoxification of heavy metals (e.g. Harada et al., 2010; Yaashikaa et al.,.2022) and may sense vibrations and mechanical signals (Zhou et al., 2017; Liu et al., 2017). Development of the branched unicellular *A. thaliana* trichomes is a multi-stage process (Mathur et al., 1999; Han et al., 2022) involving genome endoreduplication, initiation and expansion of trichome branches, and production of secondary cell wall with a characteristic structure and patterns of callose and metal ion deposition (Kulich et al., 2015; Kulich et al., 2018). The ablilty to immobilize metal ions in the trichome cell wall may contribute to detoxication of heavy metals such as excessive zinc (Ricachenevsky et al., 2021) or cadmium (Gao et al., 2021; Guo et al., 2022) by sequestration.

Knowledge regarding *A. thaliana* epidermal development comes mainly from studies on a handful of inbred lineages maintained for decades in the laboratory, such as the widespread Columbia accession. Nevertheless, for some epidermal traits, phenotypic variation among natural Arabidopsis accessions has been attributed to sequence polymorphism in specific genes, such as the *GL1, ETC2* and *MYC1* transcription regulators affecting trichome density (Hauser et al., 2001; Hilscher et al., 2009; Symonds et al., 2011; see also Hauser, 2014) or the *STI* gene controlling trichome branch number (Ilgenfritz et al., 2003).

With the onset of high throughput sequencing and availability of genetically characterized resources linked to computational tools, genome-wide association studies (GWAS) identifying sequence polymorphisms correlated with distinct phenotypic features became a powerful tool for discovering new gene functions. While the GWAS methodology is well established in Arabidopsis (see, e.g., Brachi et al., 2011; Slovak et al., 2015), it found only limited use in studying epidermal development in this species. A screen for host-side genetic determinants of plant-associated microbiome variability revealed genomic regions enriched in genes participating in cell wall biogenesis, and somewhat surprisingly, also in trichome branching (Horton et al., 2014). A recent report applied GWAS to identify genes responsible for variation in aerial organ trichome density (Arteaga et al., 2022). Epidermal or trichome characteristics have, however, been addressed by GWAS in non-model plant species, including commercially important crops. Recent studies focused on trichome distribution patterns in *Brassica napus* (Xuan et al., 2020) and *Aegilops tauschii* (Mahjoob et al., 2022), on cuticular wax deposition in *B. napus* (Jin et al., 2020; Long et al., 2023) and on cotton trichome development (Li et al., 2020; Wang et al., 2021).

Here we report a GWAS for genes associated with phenotypic variability of selected epidermal traits of rosette leaves in 310 sequenced *A. thaliana* accessions from the 1001 Genomes collection (1001 Genomes Consortium, 2016), focusing on traits related to trichome development, cell wall composition and epidermal metal deposition. Besides identifying genes known or already suspected to affect these characteristics, our results suggest possible roles in epidermal development for several gene families with hitherto uncharacterized or only partly characterized functions.

## Results

### Natural variation of rosette leaf epidermis traits

We analyzed 310 natural *A. thaliana* accessions, mostly with available detailed location data and climate parameters of their sites of origin (Supplementary Table S1) to gain insight into the genetic basis of variability in trichome shape, size and density, epidermal autofluorescence and epidermal metal deposition patterns. We recorded 7 continuous quantitative and 11 categorical semiquantitative (ordinate) or qualitative parameters (Table 1; Bezvoda, 2023) that exhibited readily noticeable variability among the accession and were amenable to semi-high-throughput analysis with good inter-observer replicability.

**Table 1.**
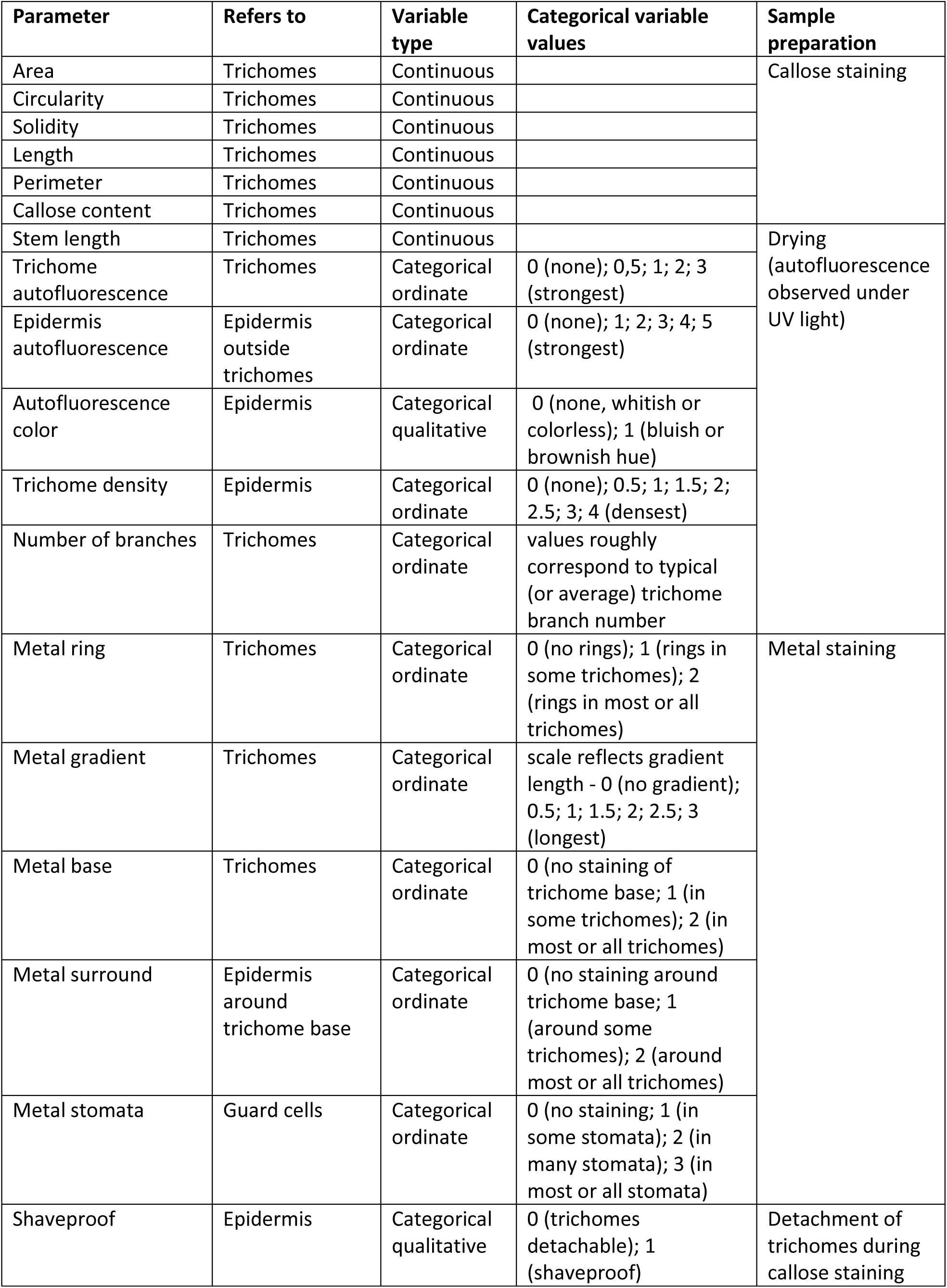
Epidermal phenotype parameters included in the GWAS screen.

| Parameter | Refers to | Variable type | Categorical variable values | Sample preparation |
| --- | --- | --- | --- | --- |
| Area | Trichomes | Continuous |  | Callose staining |
| Circularity | Trichomes | Continuous |  |  |
| Solidity | Trichomes | Continuous |  |  |
| Length | Trichomes | Continuous |  |  |
| Perimeter | Trichomes | Continuous |  |  |
| Callose content | Trichomes | Continuous |  |  |
| Stem length | Trichomes | Continuous |  | Drying (autofluorescence observed under UV light) |
| Trichome autofluorescence | Trichomes | Categorical ordinate | 0 (none); 0.5; 1; 2; 3 (strongest) |  |
| Epidermis autofluorescence | Epidermis outside trichomes | Categorical ordinate | 0 (none); 1; 2; 3; 4; 5 (strongest) |  |
| Autofluorescence color | Epidermis | Categorical qualitative | 0 (none, whitish or colorless); 1 (bluish or brownish hue) |  |
| Trichome density | Epidermis | Categorical ordinate | 0 (none); 0.5; 1; 1.5; 2; 2.5; 3; 4 (densest) |  |
| Number of branches | Trichomes | Categorical ordinate | values roughly correspond to typical (or average) trichome branch number |  |
| Metal ring | Trichomes | Categorical ordinate | 0 (no rings); 1 (rings in some trichomes); 2 (rings in most or all trichomes) | Metal staining |
| Metal gradient | Trichomes | Categorical ordinate | scale reflects gradient length - 0 (no gradient); 0.5; 1; 1.5; 2; 2.5; 3 (longest) |  |
| Metal base | Trichomes | Categorical ordinate | 0 (no staining of trichome base); 1 (in some trichomes); 2 (in most or all trichomes) |  |
| Metal surround | Epidermis around trichome base | Categorical ordinate | 0 (no staining around trichome base); 1 (around some trichomes); 2 (around most or all trichomes) |  |
| Metal stomata | Guard cells | Categorical ordinate | 0 (no staining); 1 (in some stomata); 2 (in many stomata); 3 (in most or all stomata) |  |
| Shaveproof | Epidermis | Categorical qualitative | 0 (trichomes detachable); 1 (shaveproof) | Detachment of trichomes during callose staining |

The first group of parameters with continuous distribution (Figure 1) describes quantitative aspects of trichome shape, i.e. size-related parameters (area, perimeter, overall length and stem length) and descriptors of shape complexity (circularity and solidity). Other morphological features, namely trichome density (Figure 2A, 2B) and branch number (Figure 2C), were evaluated semiquantitatively.

**Figure 1.**
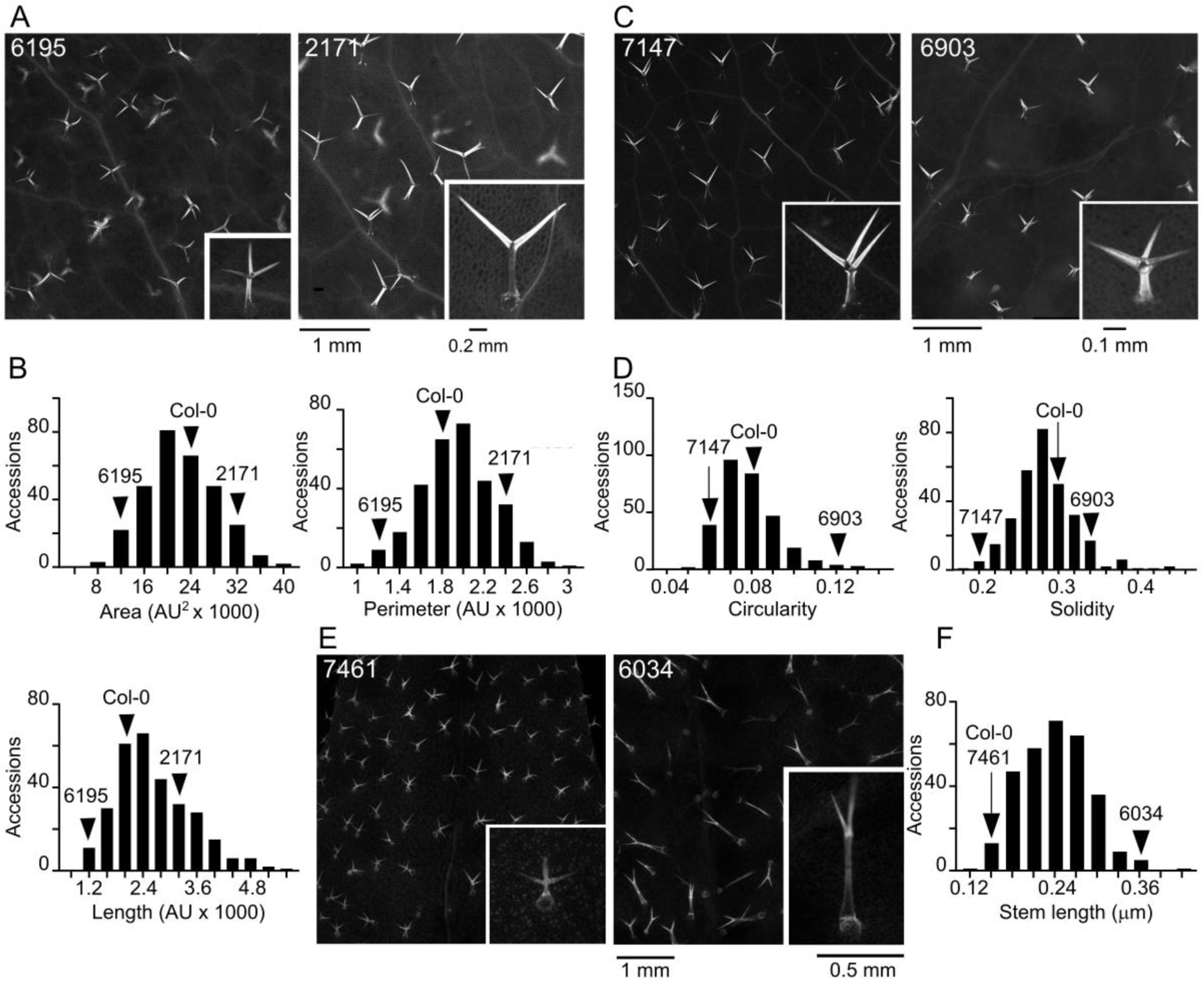
Examples of continuous quantitative parameters of trichome shape. (A) Detailed view of representative leaves of ecotypes with low (6195, TDr-9) and high (2171, Paw-26) trichome area, perimeter and length with close-up of a typical trichome. (B) Histograms showing distribution of area, perimeter and length among analyzed accessions. (C) Detailed view of representative leaves of ecotypes with low (7147, Gie-0) and high (6903, Bor-4) trichome circularity and solidity with close-up of a typical trichome. (D) Histograms showing distribution of circularity and solidity among analyzed accessions. (E) Detailed view of representative leaves of ecotypes with low (7461, H55) and high (6034, Hov1-7) trichome stem length with close-up of a typical trichome. (F) Histogram presenting distribution of stem length among analyzed accessions. Positions of listed accessions and Col-0 in the histograms are marked by arrowheads.AU - arbitrary units.

**Figure 2.**
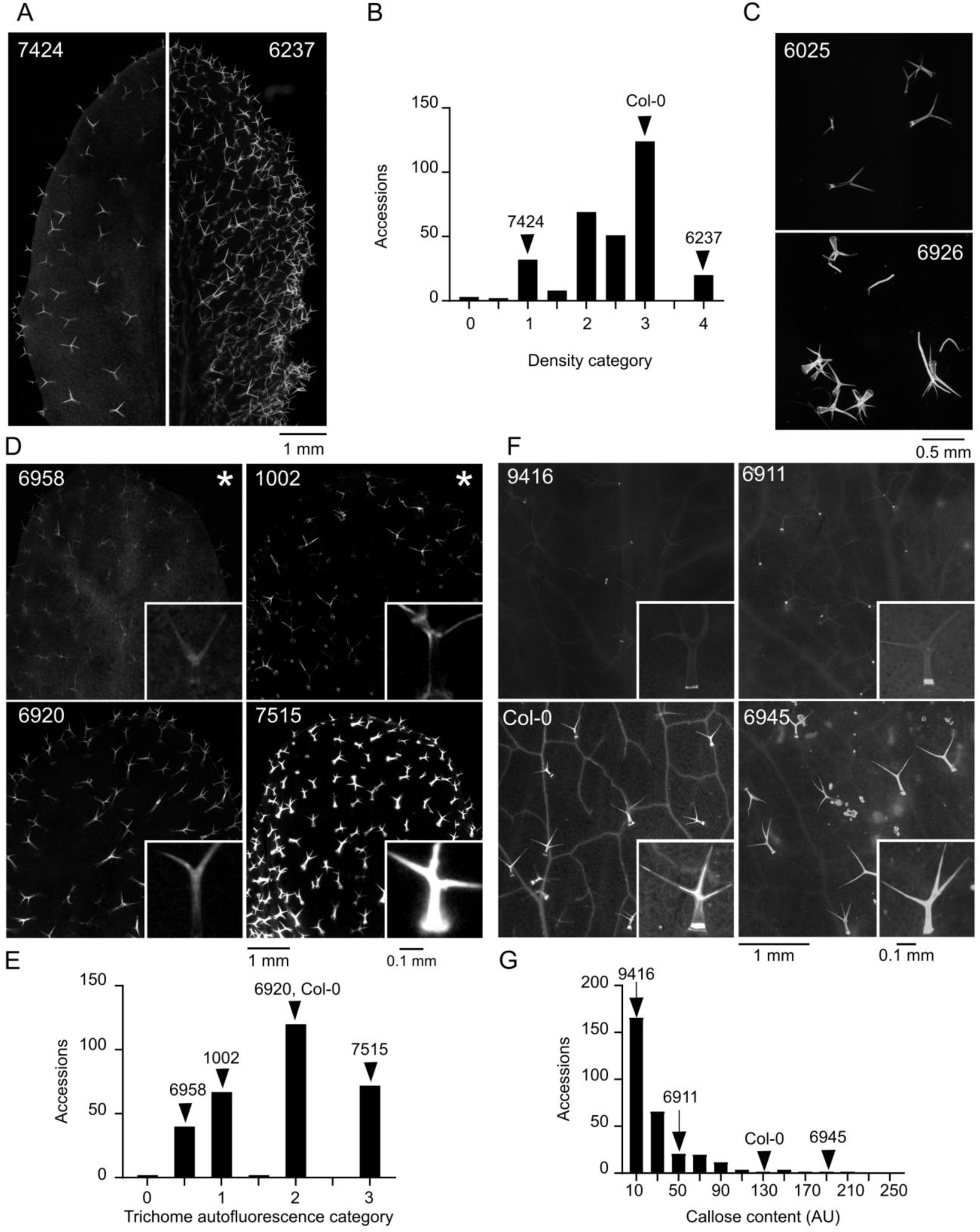
Variability of trichome density, branch number, autofluorescence and callose content. (A) Representative leaf photos of the indicated ecotypes with low (7424, JI-3) and high (6237, TOM 03) trichome density. (B) Histogram showing distribution of trichome density categories. (C) Typical isolated trichomes of an accession with low (6025, Gro-3) and high (6926, Kin-0) branch numbers. (D) Detailed view of representative leaves of ecotypes with different levels of trichome cell wall autofluorescence. (6958, Ra-0; 1002, Ale-Stenar-64-24; 6920, Got-22; 7515, RRS-10), with close-ups of single typical trichomes. Ecotypes marked by asterisks also represent typical examples of low/absent (1002, Ale-Stenar-64-24) and high (6958, Ra-0) epidermal autofluorescence outside trichomes. (E) Histogram showing distribution of trichome autofluorescence categories. (F) Detailed view of representative leaves of ecotypes with different levels of callose deposition in trichomes (9416, Ktu-3; 6911, Cvi-0; 6909, Col-0; 6945, Nok-3), with close-ups of a single typical trichome. (G) Histogram showing distribution of callose content. Positions of listed accessions and Col-0 in the histograms are marked by arrowheads.

The next group of parameters reflects structural and chemical properties of the trichome cell walls, affecting trichome autofluorescence upon damage, estimated as a semiquantitative categorical variable (Figure 2D, 2E) and callose content, evaluated semiquantitatively on a continuous scale in a manner capturing also the spatial distribution of callose (Figure 2F, 2G). Obvious variability in the background epidermal autofluorescence outside trichomes was recorded as a categorical semiquantitative variable (Figure 2D). We introduced another categorical qualitative variable to describe visible differences in the color of trichome and background fluorescence (Figure 3A).

**Figure 3.**
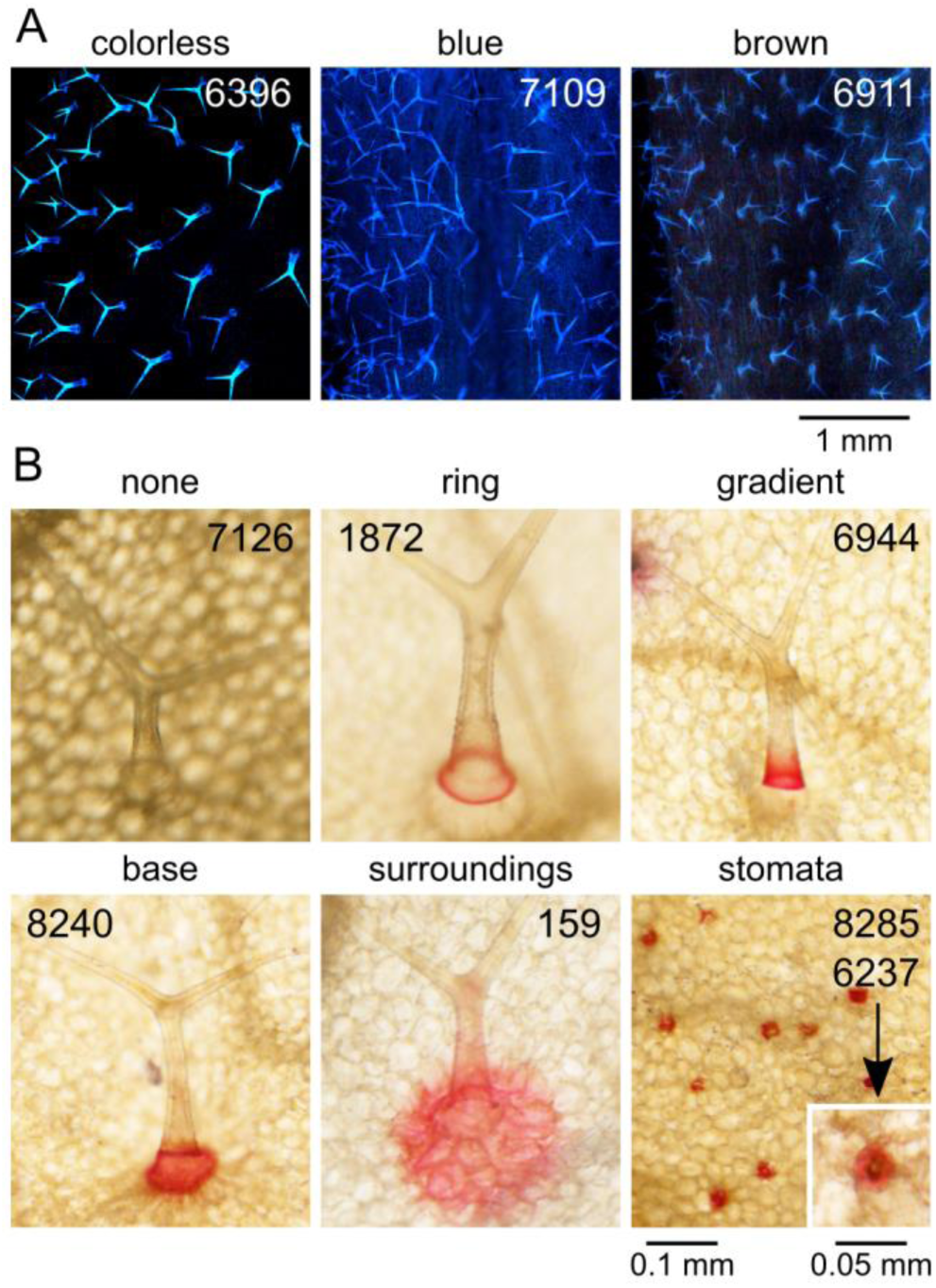
Examples of autofluorescence colors and patterns of trichome and epidermal metal deposition. (A) Representative examples of epidermal autofluorescence colors recorded as “colorless” (no background fluorescence, whitish trichomes - 6396, Udul 4-9) and “colored” (blue - 7109, Ema-1; brownish - 6911, Cvi-0). Contrast of the photographs has been uniformly adjusted using ImageJ’s “auto” settings. (B) Images of representative trichomes or epidermis areas for each pattern of metal deposition observed in the screen. Presented pattern is not necessarily the most typical or the only one found for the ecotype shown (none - 7126, Es-0; ring - 1872, MNF-Pot-75; gradient - 6944, NFA-8; base - 8240, Kulturen-1; surround - 159, MAR2-3; stomata - 8285, DraIII-1; stomata close-up - 6237, TOM 03).

A third group of categorical semiquantitative parameters captures the spatial distribution of metal ion deposits, as detected by dithizone staining, in trichomes and other epidermal cell types (Figure 3B; for description of parameters see Table 1).

Since we noticed that trichomes of some accessions resist mechanical detachment during staining for callose, we recorded this feature as a qualitative parameter “shaveproof”.

All seven continuous variable parameters exhibit smooth distribution with a single roughly symmetrical maximum (Figure 1B, 1D, 1F; Figure 2G; Figure 4A), except callose content, where most accessions have very little if any callose in trichomes, resulting in an asymmetric distribution. A roughly symmetrical single maximum distribution, in some cases noisy, was observed also for the semiquantitative categorical variables of trichome autofluorescence, epidermis autofluorescence, trichome density and trichome branch number. The “noisy single maximum distribution” may be explainable by under-representation of visually attributed intermediate values, perhaps reflecting observer uncertainty rather than genuine differences. For several other parameters (metal gradient, metal surround, metal stomata) the distribution was asymmetric with most accessions exhibiting low or no signal. Remaining parameters either varied between only two values (autofluorescence color, shaveproof) or showed a possibly bimodal distribution, such as in the case of metals at trichome base and metal rings (Supplementary Figure S1).

**Figure 4.**
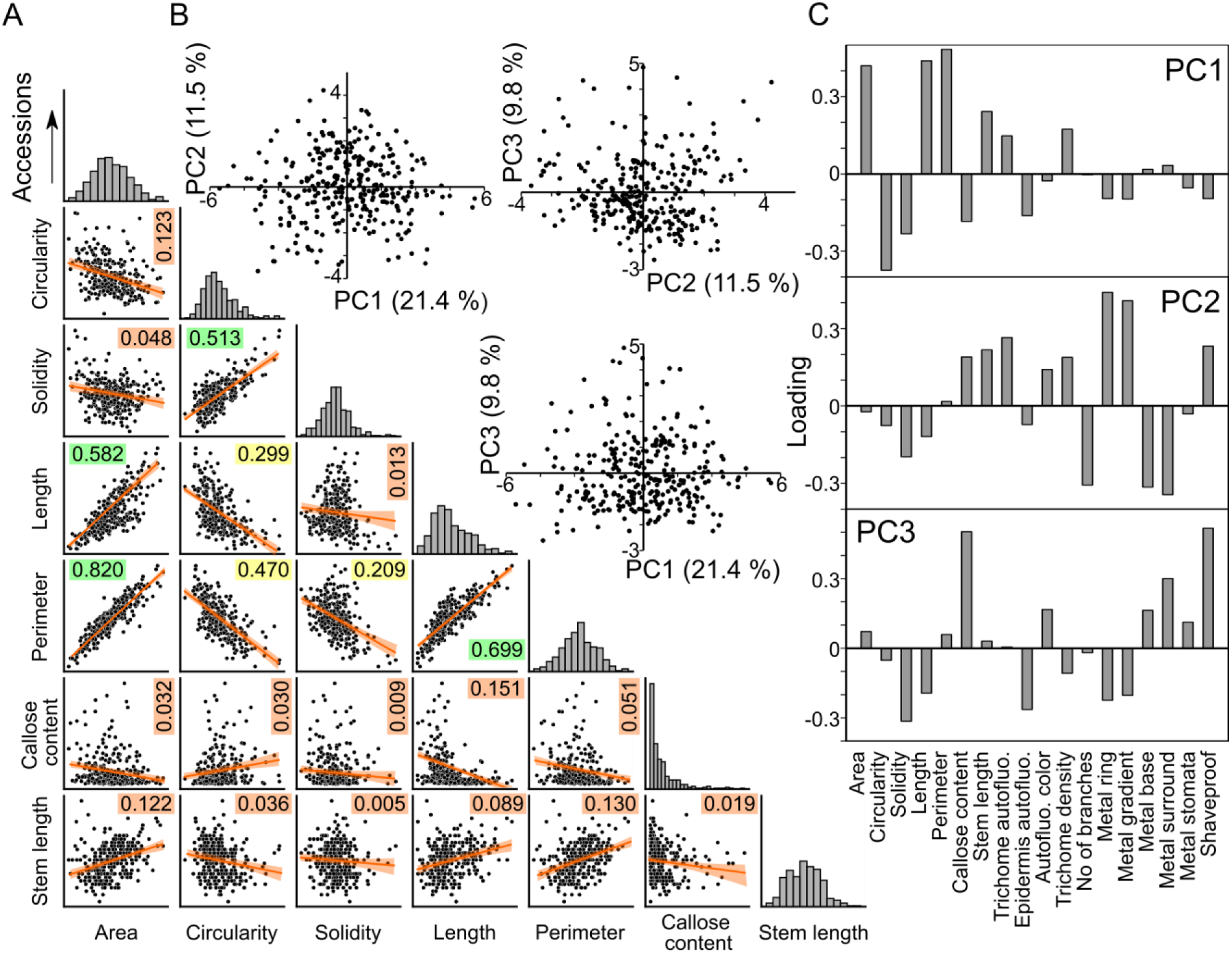
Distribution of epidermal phenotypic variability among accessions. (A) Mutual correlation of the 7 continuous-value quantitative parameters analyzed, with regression lines, confidence intervals and coefficient of determination (R^2^) values indicated. R^2^ values are color-coded from green (high, above 0.5) through yellow (intermediate) to orange (below 0.2). For Spearman’s correlation coefficients and P-values see Supplementary Figure S1. Distribution of values is shown for each parameter at the diagonal of the matrix (scales are arbitrary, for quantitative values see Figure 1B, 1D, 1F and Figure 2G). (B) Scatterplot of PCA results reflecting the between-accession variability, calculated from the complete epidermal phenotype dataset (points correspond to individual accessions). Fraction of variance attributable to each PC is shown in parentheses. (C) Loadings of the first three principal components.

Not surprisingly, some quantitative descriptors of trichome shape were mutually correlated (Figure 4A, Supplementary Figure S1). Trichome area, perimeter and overall length, which can all be viewed as measures of trichome size, exhibited strong or very strong positive correlation. Weak to moderate positive correlation with these parameters was found also for trichome stem length. Circularity (proportional to the ratio of an object’s area and squared perimeter) and solidity (ratio of the area of the examined object and its convex hull), which both achieve higher values for less intricate shapes, were mutually moderately positively correlated, and negatively correlated with the size-related parameters. While there were some cases of non-negligible correlation among the categorical variables, or between categorical and continuous ones, such correlations were generally weak except of moderate positive correlation between the frequency of metal rings and intensity of metal gradients at the trichome stems, as well as moderate positive correlation between strength of mechanical attachment (the “shaveproof” trait) and trichome callose content, which was also moderately negatively correlated with trichome length. Remarkably, all the strong or moderate correlations were statistically significant (Supplementary Figure S1).

For a subset of traits including representatives of the mutually correlated trait groups, heritability was estimated from data collected separately for individual plants (Supplementary Table S2). The broad sense heritability values, ranging between 0.58 and 0.94, indicate a strong genetic contribution to the observed variability (Supplementary Table S3).

Principal component analysis (PCA) of the 18 recorded parameters did not reveal any obvious clustering of accessions when considering the first three principal components (PCs), which, however, cumulatively explained only less than 43 % of total variability (Figure 4B). Analysis of PC loadings indicated that PC1 is positively correlated with the mutually correlated parameters reflecting trichome size, and negatively with circularity and solidity, PC2 correlates either positively or negatively with specific patterns of metal staining, and PC3 mostly reflects callose content and strength of mechanical attachment - i.e. two parameters related to cell wall structure (Figure 4C). The PCA results thus support the overall pattern of continuous variability of traits addressed in our study, and also suggest that trichome shape, metal staining and cell wall organization might be determined by distinct sets of genes.

We next examined possible relationships between the 18 phenotypic traits determined in our study and 35 standardized climate variables of the sites of origin of our Arabidopsis accessions, obtained from the CliMond database (Kriticos et al., 2012). Out of the 630 possible trait and climatic variable combinations, only 81 exhibited statistically significant non-negligible positive or negative correlation, typically in the weak range (Supplementary table S4, Figure 5A). The only exception was a negative correlation between trichome stem length and the maximum temperature of the warmest week (the Bio05 variable), which fell into the moderate range (*sensu* Schober et al., 2018; Figure 5B). The biological interpretation of this observation is not immediately obvious.

**Figure 5.**
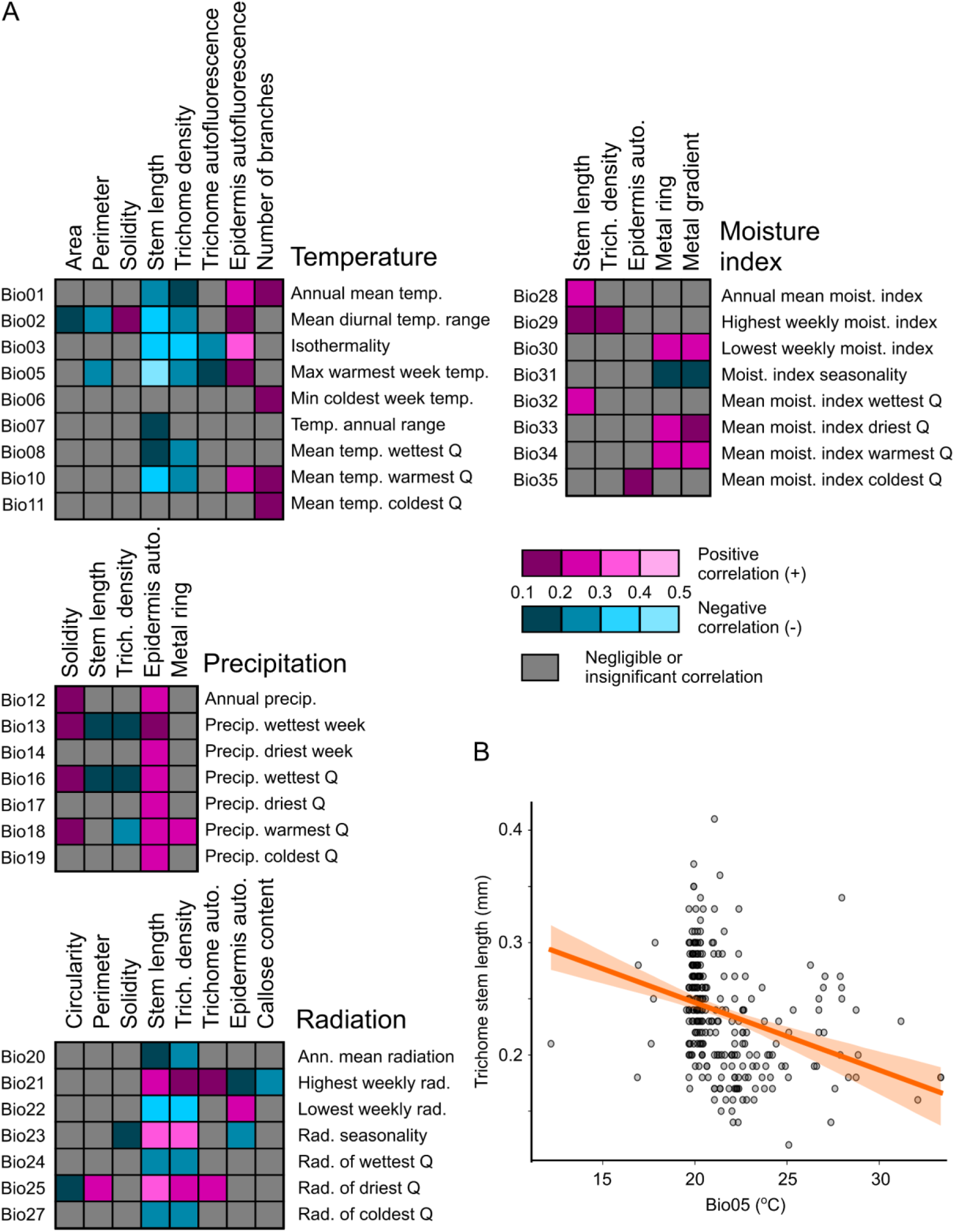
Correlations of epidermal traits and climatic variables. (A) Heatmaps summarizing all statistically significant non-negligible correlations among epidermal traits and BioClim variables of the site of origin, grouped according to underlying environmental factors and color-coded according to the Spearman’s correlation coefficient values. For a complete list of combinations, correlation coefficient values and P-values see Supplementary Table S4. (B) Correlation of trichome stem length and the maximum temperature of the warmest week at the site of origin (variable Bio05). The coefficient of determination (R^2^) value was 0.204, i.e. in the intermediate range.

### GWAS analysis identifies loci associated with some epidermal traits

The phenotypic data were subjected to association analysis using the publicly available GWAPP GWAS platform (Seren et al., 2012) as described in Materials and Methods, aiming first towards identification of all single nucleotide polymorphisms (SNPs) within the transcribed section of genomic DNA, including the CDS, introns and the 5’ and 3’ UTRs. The numbers of identified significantly associated loci varied among traits on the scale from zero to several hundred, with trichome stem length and guard cell metal accumulation yielding the most candidates (Table 2; Supplementary Table S5). No significant SNPs were found for trichome area, metal ring, metal staining of trichome bases, and resistance towards mechanical detachment, and only one or a few presumably silent polymorphisms (i.e. synonymous mutations or SNPs located in an intron or in untranslated mRNA regions) were identified for overall trichome length, number of branches, and the presence of a metal gradient. These traits were therefore not included in subsequent analyses.

**Table 2.**
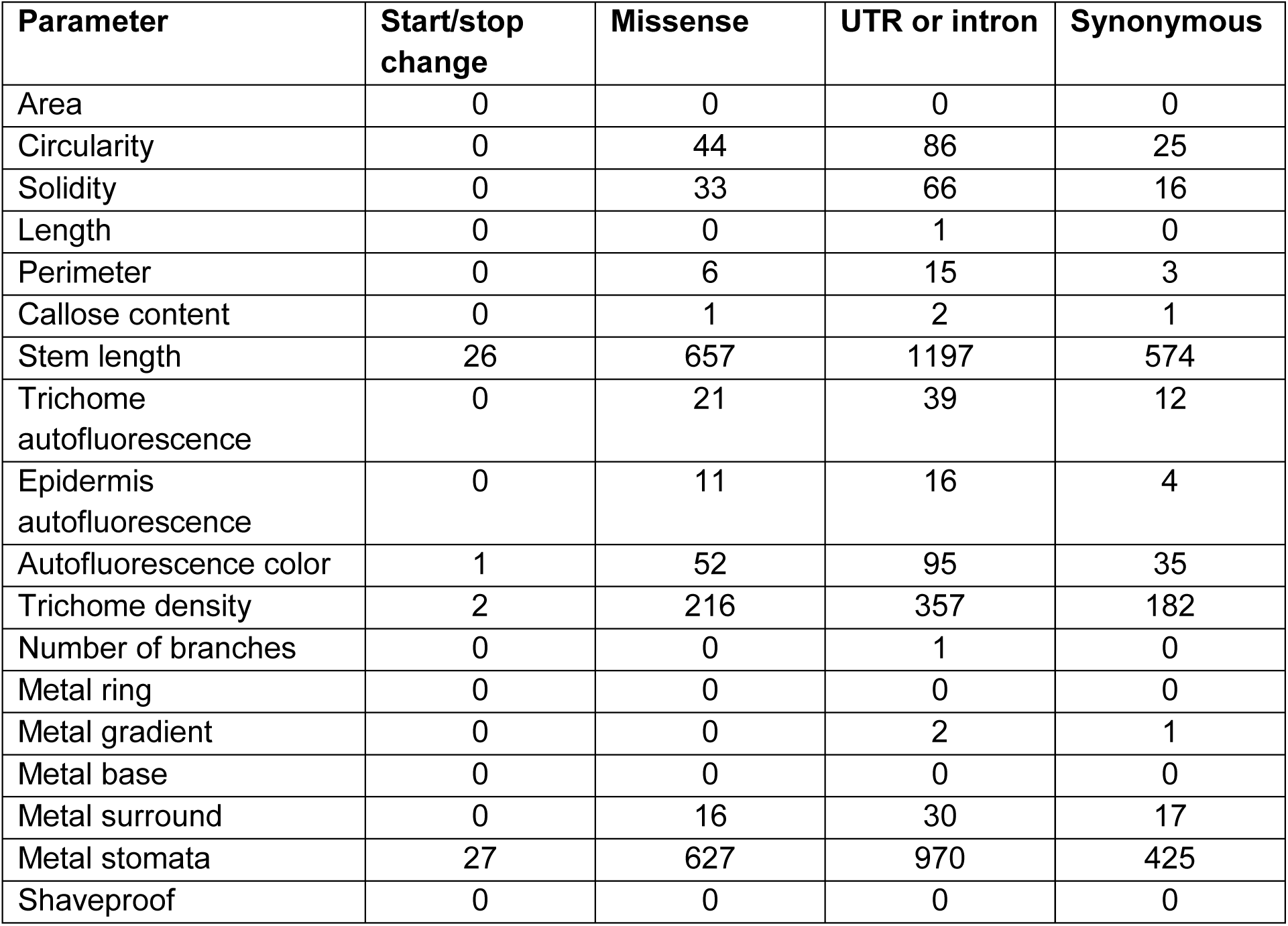
Numbers of loci and polymorphism types identified as significantly associated with individual parameters.

The distribution of SNP types for individual traits with multiple SNPs exhibits some differences, although they are not major (Supplementary Figure S2). In particular, SNPs altering CDS length were found only for traits with overall large SNP numbers, reflecting low probability of polymorphisms affecting start and stop codons. Subsequently, we focused only on loci with SNPs changing the predicted protein product sequence, in most cases due to a missense mutation, and treated all such SNPs as a single category. The full list of genes affected by such SNPs for each trait is provided in Supplementary Table S6. In total, 1547 different loci exhibited association with at least one of the studied traits, corresponding to approximately 5.6 % of all Arabidopsis genes.

For further analyses, we divided the traits with significantly associated SNPs into four groups: those affecting trichome shape (circularity, solidity, perimeter and stem length), trichome distribution on the surface of the leaf (i.e. trichome density), wall composition-related parameters (epidermis and trichome autofluorescence, autofluorescence color, callose content) and metal accumulation (metal staining of stomata and trichome surroundings). While some loci associated with several of the mutually correlated cell shape traits, there was only a very limited, if any, overlap between candidate lists for traits from the wall composition-related and metal accumulation-related groups. However, polymorphisms in some loci were identified as associating with traits from more than one category (Figure 6A, Supplementary Table S7).

**Figure 6.**
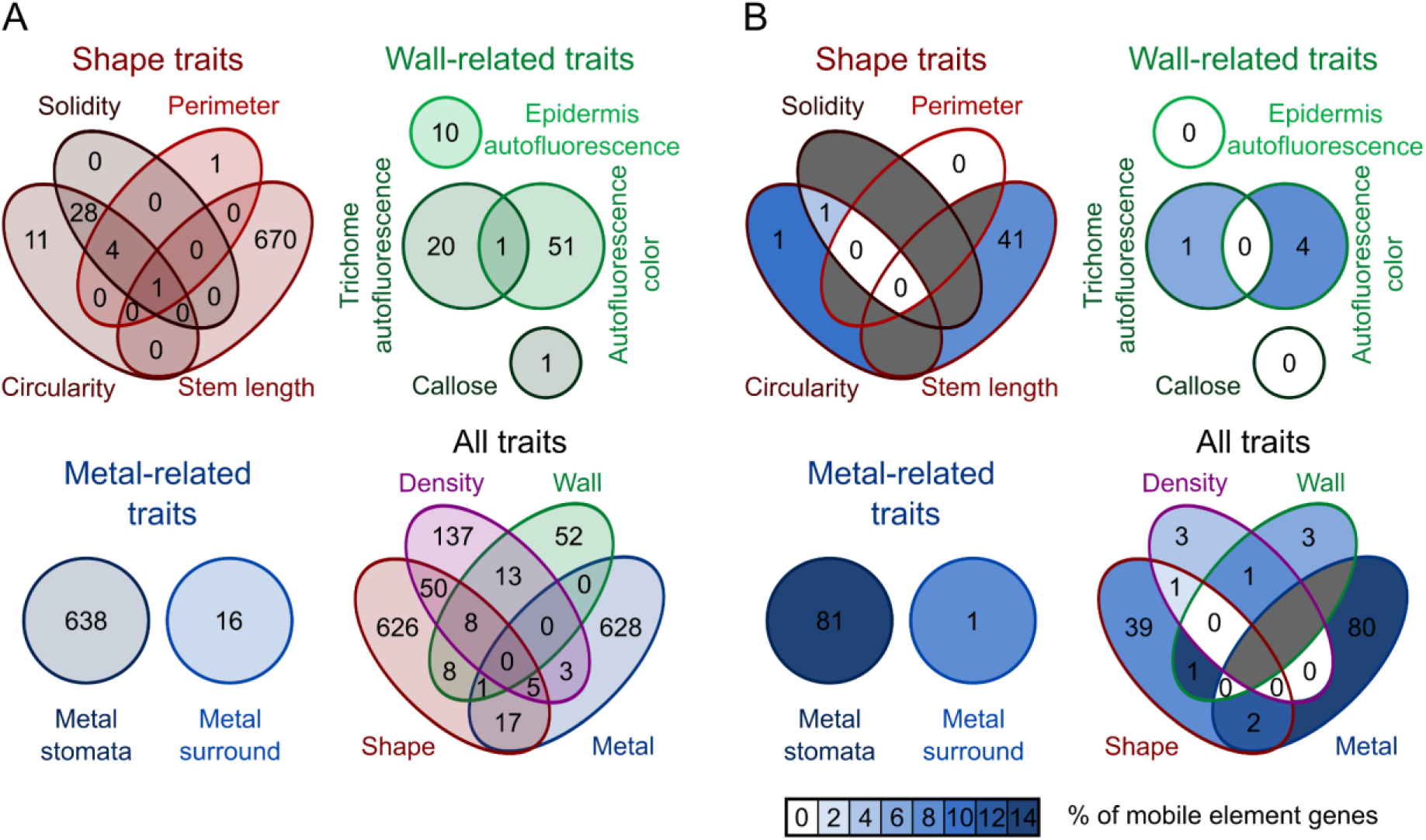
Unique and shared SNPs among traits and trait groups. Venn diagrams showing numbers of loci affected by ORF-changing SNPs associated with individual traits and trait groups and/or shared by multiple traits or trait groups. (A) All ORF-changing SNPs.. (B) ORF-changing SNPs in loci contained within transposons. Fields are color-coded according to proportion of transposon-derived loci among all loci detected for the given trait combination. Dark gray fields indicate trait combinations with no associated SNPs detected.

Assuming that SNPs in transposon-borne genes are unlikely to cause phenotypic differences, we examined what fraction of our candidates corresponds to CDSs from mobile genetic elements to gain insights into the extent of non-specific background. The fraction of transposon-derived loci was significantly depleted compared to whole genome values for the sum of all GWAS candidates, as well as for those associated with shape-related traits or trichome density. However, while depletion of transposon genes was also noticeable for wall-related traits, it was only marginally significant, possibly due to a relatively low number of loci, and practically no depletion was observed for metal-related traits (Table 3). Sets of genes associated with multiple loci usually contained fewer transposon-derived loci than those found on the basis of a single trait (Figure 6B).

**Table 3.** Distribution of mobile element-derived genes among candidate loci associated with ORF-changing SNPs.

| Trait group | All loci | Transposon loci | Fraction from transposons | P-value |
| --- | --- | --- | --- | --- |
| Whole genome (Araport) | 27655 | 3879 | 14 % | N.A. |
| All traits | 1547 | 129 | 8 % | <0 .0001 |
| Shape traits | 715 | 43 | 6 % | <0 .0001 |
| Density | 216 | 5 | 2 % | <0 .0001 |
| Wall-related traits | 82 | 6 | 7 % | 0.10 |
| Metal-related traits | 654 | 82 | 13 % | 0.25 |

P-values indicate significance of the transposon-derived locus depletion compared to the whole genome dataset (pairwise Chi-square test with Benjamini-Hochberg correction for multiplicity). N.A. - not available.

### Multiple GWAS candidates encode proteins with known or suspected epidermal roles

Next, we examined the traits related to trichome development (i.e. the shape traits, density, and the cell wall parameters that largely reflect trichome properties) for enrichment of genes abundantly expressed in trichomes either at the mRNA (Jakoby et al., 2008) or protein (Huebbers et al., 2022) levels. While we did not detect significant enrichment or depletion of these genes in our candidates lists, we identified multiple genes associated with trichome related traits that were abundantly expressed in trichomes, consistent with their participation in trichome development. In some cases, available annotations suggest function in membrane trafficking, biogenesis of cell wall components or turgor generation and maintenance. This might point to possible mechanisms whereby variation in the responsible loci may affect cell growth or cell wall development and generate the observed phenotypic diversity, either by direct modification of trichome morphogenesis or by modulating epidermal cell expansion and hence trichome density (Table 4). Prompted by the finding of several genes engaged in cuticle or epidermal wax biosynthesis, i.e. traits often linked to trichome development (see Berhin et al., 2022) among these loci, we examined the rest of our candidates for overlap with a list of 87 putative cuticle and wax biosynthesis genes identified on the basis of their known activities and/or expression patterns (Suh et al., 2005). However, we did not find any additional loci in this manner beyond two genes that were also highly expressed in trichomes - namely AT1G01600/CYP86A4 and AT1G76690/OPR2 (see Table 4).

**Table 4.** Candidate loci associated with trichome development traits and highly expressed in trichomes, with phenotypic effects of identified substitutions indicated.

| Locus (Gene name) | Screen | Description | Trait | Allele (Effect) |
| --- | --- | --- | --- | --- |
| AT1G01600 (CYP86A4) | T | Member of the CYP86A subfamily of cytochrome p450 genes. Involved in cutin and wax biogenesis (Camoirano et al., 2020). | Stem length | V341I (up) |
|  |  |  | Density | A476V (down) |
|  |  |  | Trichome autofluorescence | A476V (down) |
| AT1G08920 (ESL1) | T | ERD (early response to dehydration) six-like 1; ESL1, a transporter for monosaccharides. | Stem length | L118F (up) |
|  |  |  | Density | L118F (up) |
| AT1G12550 (HPR3) | P | Glyoxylate/hydroxypyruvate reductase HPR3. Transcript restricted to trichomes (Xu et al., 2018). | Stem length | S135A (up) |
| AT1G12570 (HTH-like) | P | Glucose-methanol-choline (GMC) oxidoreductase family protein. Ortholog of maize IPE1 participating in pollen exine development. Involved in cutin and wax biogenesis (Kannangara et al., 2007). | Stem length** | K138R (up)<br>N187K (up) |
|  |  |  | Density | D77G*, S78A* (up) |
| AT1G15670 (KFB1, KMD2) | T | F-box/Kelch-repeat protein. Member of the KISS ME DEADLY (KMD) family, that targets type-B ARR proteins for degradation and is involved in negative regulation of the cytokinin response. Also known as KFB1, a member of a group of Kelch repeat F-box proteins negatively regulating phenylpropanoid biosynthesis. | Stem length | E37A (up) |
| AT1G26400 (FOX3, ATBBE5) | P | Berberine bridge enzyme-like 5. Possible role in defense, related proteins participate in wall modification (Eggers et al., 2021). | Stem length | I16V*, L39P* (up) |
| AT1G53210 (NCL) | P | Sodium/calcium exchanger NCL. | Stem length | G420D (down) |
| AT1G76690 (OPR2) | P | 12-oxophytodienoic acid reductase with possible role in cutin or wax synthesis (Suh et al., 2005). | Stem length | A90T (down) |
| AT1G78920 (AVPL1) | P | Pyrophosphate-energized membrane proton pump 2. Located at Golgi, strong expression in trichomes confirmed using a GUS reporter (Mitsuda et al., 2001). | Stem length | N16S (up) |
| AT2G20240 (TRM17) | P | GPI-anchored adhesin-like protein, putative (DUF3741). | Stem length | S157P, D576E*, A671T* (up) |
| AT2G20990 (SYTA, SYT1) | P | Synaptotagmin A. Regulates endosome recycling at the plasma membrane, but not secretory traffic. Strong expression in trichomes confirmed using a GUS reporter (Schapire et al., 2008). | Stem length | H357Q (up) |
| AT2G21410 (VHA-A2) | P | V-type proton ATPase subunit a2. Required for endocytic and secretory trafficking (Dettmer et al., 2006). | Stem length | I519V*, I762V (up) |
| AT2G21870 (MGP1, PHI1) | P | Probable ATP synthase 24 kDa subunit, mitochondrial. Essential for pollen formation. | Stem length | K207M (up) |
| AT2G35860 (FLA16) | P | Fasciclin-like arabinogalactan protein. Mutant has reduced cell size in some cell types in the stem (Liu et al., 2020). | Density | K115T (down) |
| AT3G28500 (RPP2C, P2X) | P | 60S acidic ribosomal protein P2-3. | Density | G82A (down) |
| AT3G48000 (ALDH2B4) | P | Aldehyde dehydrogenase family 2 member B4, mitochondrial. May participate in biosynthesis of ferulic acid and sinapic acid, modulating cell wall strength (Brocker et al., 2013). | Stem length | N63D (up) |
| AT3G59010 (PME35, PME61) | P | Pectin methyl esterase, activity controls cell wall mechanic strength; identified in a trichome transcriptome study (Wienkoop et al., 2004) | Density | V291I (down) |
| AT4G02930 (TUFA) | P | Elongation factor Tu, mitochondrial. | Stem length | V29I (down) |
| AT4G24840 (COG2) | P | Oligomeric Golgi complex subunit-like protein containing the COG2 domain, part of the membrane trafficking machinery (see Vukašinović and Žárský, 2016) | Stem length | L652F (down) |
| AT5G28220 | P | Protein prenyltransferase (uncharacterized). | Stem length | L190F (down) |
| AT5G63910<br>(FCLY) | P | Farnesylcysteine lyase (EC 1.8.3.5) involved in a salvage /detoxification of farnesylcysteine residues liberated during the degradation of prenylated proteins (Crowell et al., 2007). | Density | G88S (down) |

Unless additional references are provided, annotation is derived from Araport 11 or the TAIR description (Reiser et al., 2024). T – transcriptome, P – proteome. Substitutions and effect directions refer to the Col-0 sequence and phenotype. \* – not documented in some of the ecotypes carrying the other listed substitution because of sequencing gaps; \*\* – the listed minor alleles affecting this parameter are often present together.

Because the loci associated with metal accumulation-related traits mainly reflect stomatal metal deposition, we examined the metal-related candidate list for overlap with a published set of 63 genes upregulated in guard cells (Leonhardt et al., 2004), as well as a list of 1508 proteins repeatedly identified in guard cell proteome (Zhao et al., 2008). We found no overlap between our candidate list and the list of genes highly transcribed in guard cells, but identified 21 loci from the proteome set, all of them associated with metal accumulation in the guard cells and some encoding proteins engaged in ion or other solute transport, membrane trafficking, cell wall biogenesis or gene expression, i.e. processes possibly relevant for the observed trait (Table 5). However, there was no significant enrichment of guard cell-expressed genes among our candidates, similar to the trichome case.

**Table 5.** Candidate loci associated with stomatal metal accumulation and expressed in the guard cell proteome.

| **Locus**<br>**(Gene**<br>**name)** | **Description** | **Allele** |
| --- | --- | --- |
| AT1G31850 | S-adenosyl-L-methionine-dependent methyltransferases superfamily protein; probable methyltransferase PMT20; found in a Golgi complex proteome study (Parsons et al., 2012) | V113A |
| AT1G42550<br>(PMI1) | Protein PLASTID MOVEMENT IMPAIRED 1, a plant-specific protein of unknown function conserved among angiosperms | G460A |
| AT1G59820<br>(ALA3) | Phospholipid-transporting ATPase (phospholipid translocase) involved in secretory vesicle formation from trans-Golgi. | E647K |
| AT1G59900<br>(E1 ALPHA) | Pyruvate dehydrogenase E1 component subunit alpha-1, mitochondrial. | S361P |
| AT1G67490<br>(GCS1, KNF,<br>KNOPF) | Mannosyl-oligosaccharide glucosidase GCS1; an alpha-glucosidase I enzyme that catalyzes the first step in N-linked glycan processing. Localized to the endoplasmic reticulum. | D567E |
| AT3G10670<br>(ABCI6,<br>NAP7) | ABC transporter I family member 6, chloroplastic. Plastidic SufC-like ATP-binding cassette/ATPase essential for Arabidopsis embryogenesis. Involved in the biogenesis and/or repair of oxidatively damaged Fe–S clusters. | L42V |
| AT3G13460<br>(ECT2) | Evolutionarily conserved C-terminal region 1; regulates the mRNA levels of the proteasome regulator PTRE1 and of several 20S proteasome subunits, resulting in enhanced 26S proteasome activity. | T210P |
| AT3G13772<br>(TMN7) | Transmembrane 9 superfamily member, localized in the secretory pathway. Overexpression of this protein in yeast alters copper and zinc homeostasis (Hegelund et al., 2010). | N565S |
| AT3G48000<br>(ALDH2B4,<br>ALDH2,<br>ALDH2A) | Aldehyde dehydrogenase family 2 member B4, mitochondrial. | A267T |
| AT4G13360 | 3-hydroxyisobutyryl-CoA hydrolase-like protein 3, mitochondrial; ATP-dependent caseinolytic (Clp) protease/crotonase family protein. | T395A |
| AT4G18290<br>(KAT2) | Member of the Shaker family potassium ion (K<sup>+</sup>) channel. Critical to stomatal opening induced by blue light and to circadian rhythm of stomatal opening. Involved in plant development in response to high light intensity. | L170H |
| AT4G18670<br>(LRX5) | Leucine-rich repeat extensin-like protein 5, involved in cell wall biogenesis and organization. Interacts with several members of the RALF family of ligand peptides. | P507T<br>P813L |
| AT4G20400<br>(JMJ14) | Histone H3K4 demethylase repressing floral transition. | P480L |
| AT4G21180<br>(ERDJ2B) | DnaJ domain protein localized in ER membrane. | T67A |
| AT5G35430<br>(NOT10) | Tetratricopeptide repeat (TPR)-like superfamily protein, found also in stress granules. | W680C |
| AT5G39040<br>(ABCB27,<br>ALS1, TAP2) | Aluminum sensitive 1, ABC transporter B family member 27; a member of TAP subfamily of ABC transporters that is located in the vacuole. Mutants are hypersensitive to aluminum. May be important for intracellular movement of some substrate, possibly chelated Al, as part of a mechanism of aluminum sequestration. | V266I |
| AT5G43130<br>(TAF4B) | Transcription initiation factor TFIID subunit 4b. | Q120K |
| AT5G52470<br>(MED36B,<br>FBR1, FIB1,<br>SKIP7) | Probable mediator of RNA polymerase II transcription subunit 36b; encodes a fibrillarin, a key nucleolar protein in eukaryotes which associates with box C/D small nucleolar RNAs (snoRNAs) directing 2'-O-ribose methylation of the rRNA. | L133V |
| AT5G55040<br>(BRD13) | DNA-binding bromodomain-containing protein; interacts with core SWI/SNF complex components. Identified as a specific subunit of BRM-associated SWI/SNF (BAS) complexes | S256A |
| AT5G58100<br>(IEF7) | Encodes a membrane protein involved in pollen nexine and sexine development. | N358H |
| AT5G58140<br>(PHOT2) | Phototropin-2; membrane-bound protein serine/threonine kinase that functions as blue light photoreceptor in redundancy with PHO1. Involved in stomatal opening, chloroplast movement and phototropism. | R898G |

Unless additional references are provided, annotation is derived from Araport 11 or the TAIR description (Reiser et al., 2024). Substitutions refer to the Col-0 sequence; all listed alleles exhibited increased guard cell metal deposition compared to Col-0.

To further verify the performance of our GWAS screen, we inspected the candidate gene lists for genes previously reported to affect the studied traits. While such a search cannot be exhaustive, we found at least five previously characterized relevant genes (Table 6), including loci coding for two cytoskeletal proteins CLASP and SCAR/DIS3, as well as the EXO84B gene, encoding a subunit of the exocyst complex that regulates targeting of membrane vesicles. All three genes exhibit mutant phenotypes involving trichome shape alterations, and all associated with trichome shape traits. In addition, the gene for another exocyst subunit, EXO70H4, documented to participate in the deposition of metals in the trichome cell wall, was picked up as associating with guard cell metal deposits. The fifth gene, GFS9/TT9, is known to affect cell wall autofluorescence and was found in association with trichome density and autofluorescence traits. Notably, five SNPs affecting this gene often occurred together, constituting a quintuple substitution minor allele associated with reduced trichome density and decreased autofluorescence.

**Table 6.** Candidate loci that were previously characterized as participating in the development of relevant epidermal traits, with phenotypic effects of identified substitutions indicated.

| Locus<br>(Gene name) | Description | Trait(s) | Allele (Effect) |
| --- | --- | --- | --- |
| AT2G20190<br>(CLASP) | CLIP-associated protein; a microtubule-associated protein involved in both cell division and cell expansion. It likely promotes microtubule stability. Loss of function mutants have decreased trichome branch number (Pietra et al., 2013) | Stem length | M1399L (up) |
| AT2G38440<br>(DIS3, ITB1, SCAR2, WAVE4) | Subunit of the WAVE complex, required for activation of ARP2/3 actin nucleation complex. Mutations cause defects in both the actin and microtubule cytoskeletons that result in aberrant epidermal cell expansion. Loss of function mutants exhibit a distorted trichome phenotype (Basu et al., 2005). | Circularity | I1265V (down) |
|  |  | Solidity | I1265V (down) |
|  |  | Density | I1265V (down) |
| AT3G09520<br>(EXO70H4) | A member of the EXO70 gene family of putative exocyst subunits, conserved in land plants. Involved in trichome cell wall maturation, required for deposition of heavy metals in the cell wall (Kulich et al., 2015, though guard cell localization was not specifically observed). | Metal stomata | S93N (up) |
| AT3G28430<br>(GFS9, TT9) | Peripheral membrane protein localized at the Golgi apparatus that is involved in membrane trafficking, vacuole development and flavonoid accumulation. Loss of function mutants exhibit abnormal autofluorescence, though trichomes were not specifically examined (Ichino et al., 2014). | Density | S573A, S626A, Y667H*, A670V*, D671E* (down) |
|  |  | Trichome autofluorescence. | S573A, S626A, Y667H*, A670V*, D671E* (down) |
| AT5G49830<br>(EXO84B) | Exocyst complex component 84B; involved in tethering vesicles to the plasma membrane. Interacts with Exo70H4 that controls trichome cell wall development; loss of function mutant has multiple defects including in trichome morphogenesis (Kulich et al., 2015; Fendrych et al., 2010). | Circularity | E123G (down) |
|  |  | Solidity | E123G (down) |

Substitutions and effect directions refer to the Col-0 sequence and phenotype; \* – not documented in some of the ecotypes carrying the other listed substitution because of sequencing gaps.

Additional 80 genes with previously reported roles in cytoskeletal organization, membrane trafficking, cell wall biogenesis, tissue patterning or other relevant functions were also found among the candidates, including several genes affecting aspects of epidermal development unrelated to traits addressed in our study (Supplementary Table S8).

These findings suggest that, despite the lower efficiency of the metal trait-associated gene search, as evident from the lack of transposon gene depletion, the set of candidates found in our screen contains at least a sizeable proportion of genes biologically relevant in the context of leaf epidermis differentiation.

### Some GWAS candidates exhibit environmentally correlated allele distribution

To obtain clues to the mechanisms underlying the observed correlation between trichome stem length and the maximum warmest week temperature of the accession’s location of origin (the Bio05 variable), we further examined the set of genes exhibiting SNPs associated with trichome stem length. Since the full list of over 600 stem length-associated candidates was too large for a detailed study, we focused on the 18 genes that were at the same time abundantly expressed in trichomes or encoded proteins with known roles in trichome development (see Table 4 and Table 6).

For twelve of these genes, at least 30 studied genotypes (i.e., a tenth of our accessions set) harbored the minor allele. The distribution of Bio05 values for these loci in all but one case exhibited a statistically significant difference between the major and minor alleles, indicating an environmentally correlated bias in allele distribution (Figure 7A), which might suggest natural selection against the minor alleles in hot habitats. Among the affected genes were loci encoding the microtubule-associated protein CLASP, known to have pleiotropic roles in cell expansion and cell division (Ambrose et al., 2007; Pietra et al., 2013), and SYT1, coding for a synaptotagmin engaged in processes related to stress tolerance and immunity towards pathogens (Schapire et al., 2008; Yamazaki et al., 2008; Levy et al., 2015; Kim et al., 2016). It this thus conceivable that the selection was operating on of some function(s) of these genes unrelated to epidermal morphogenesis.

**Figure 7.**
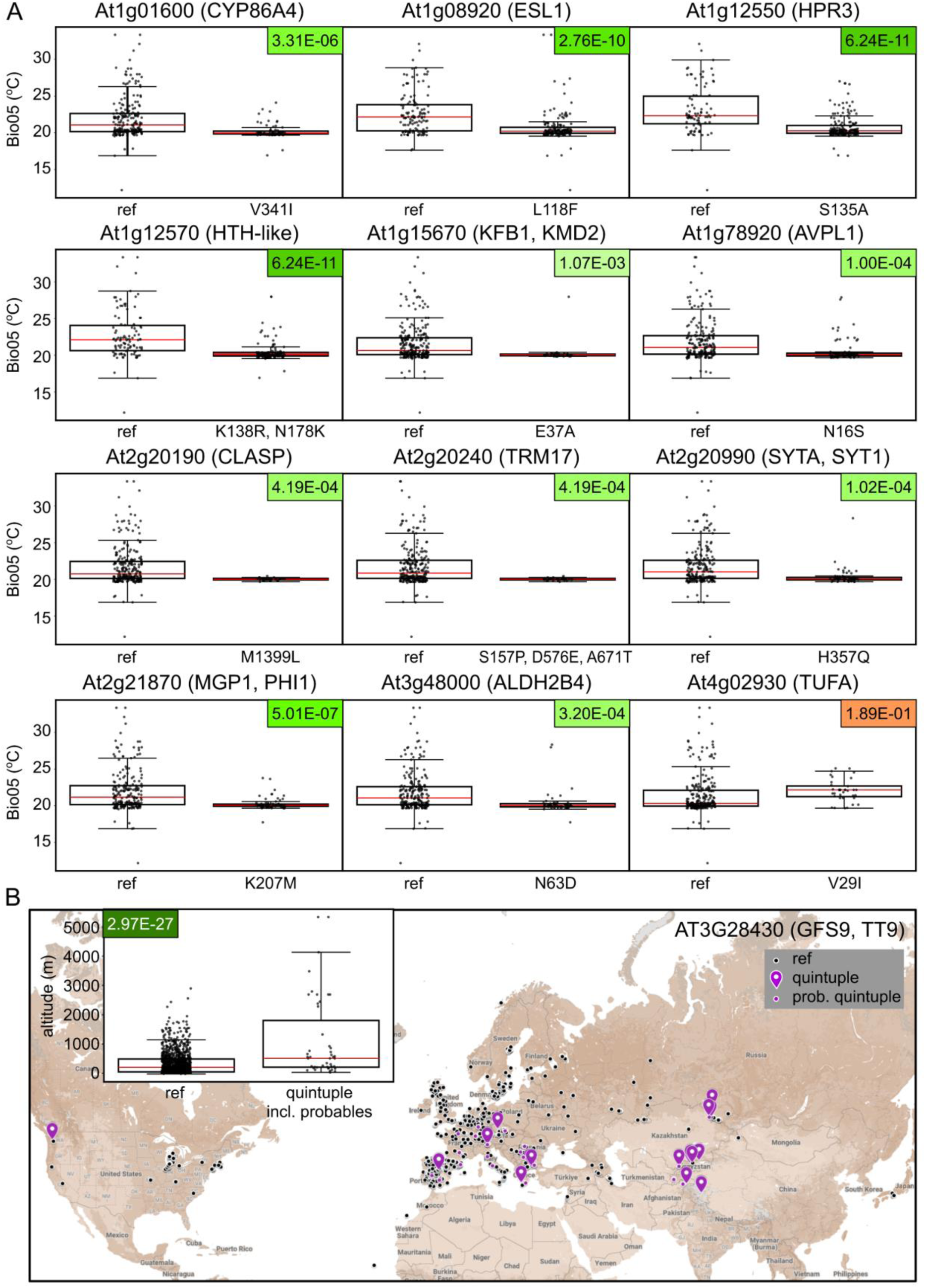
Environmentally correlated allele distribution of selected loci. (A) Distribution of the Bio05 climatic variable values for reference and minor alleles of selected genes linked to the trichome stem length trait among our experimentally characterized Arabidopsis accessions. (B) Geographic distribution of populations from the whole Ensembl data set carrying the reference vs. quintuple minor (S573A, S626A, Y667H, A670V, D671E) allele of GFS9/TT9. Accessions where some of the SNPs characteristic for the quintuple allele are missing due to sequencing gaps are categorized as “probable quintuple”. Inset – distribution of the altitude parameter among accessions with the reference or quintuple minor allele of GFS9/TT9 (including the probable quintuple substitution accessions). FDR-corrected P-values for the observed differences are color-coded green for highly significant, 0 < P ≤ 0.01 (with darker color corresponding to lower P) or orange for not significant (P > 0.05).Red lines indicate median values.

For all genes with significant environmentally correlated allele distribution, populations harboring minor alleles originated from multiple sites, i.e. there was no evidence of spatial restriction due to a founder effect. Even variants only found in Scandinavian accessions occured at two or more geographically distant sites (Supplementary Figure S3). This further supports the relevance of environmental factors and possible natural selection.

Since the GFS9/TT9 gene displayed an unusual co-occurrence of five SNPs, as described above, we also examined the geographic substitution of the quintuple substitution minor allele. Because the five contributing substitutions were found in mere six accessions of our experimentally characterized collection, we analyzed the whole set of genotypes from the Ensembl database (Yates et al., 2022). Populations carrying the quintuple SNP allele often originated from mountainous regions, and the average altitude of their sites of origin was significantly higher than for the reference variant (Figure 7B). Correspondingly, a significant difference was found in multiple climatic variables from the standard BioClim set, especially in parameters reflecting typical features of high altitude biotops, such as increased temperature seasonality, greater difference among temperature and precipitation extremes, and increased solar radiation (Supplementary Figure S4).The presence of the quintuple substitution allele across three continents is consistent with its selective advantage under high altitude conditions.

### Overrepresentation of some gene families among GWAS candidates suggests new gene functions

We next examined the sets of loci associated with the four trait groups (trichome shape, trichome density, wall-related traits and metal-related traits) for Gene Ontology term association. No enrichment or depletion of any GO categories was found on the P = 0.01 significance level, while on the P = 0.05 level, a 2.73 x enrichment of the Transmembrane signal receptor protein class was found among shape-associated loci, and a 12.22 x enrichment of the snoRNA-binding molecular function category was observed among metal trait-associated loci. The biological significance of the latter, however, remains unclear in the light of the apparent low reliability of metal-related candidate detection (see above).

While inspecting our candidate gene lists, we noticed the presence of multiple members of some gene families, including those encoding evolutionarily conserved domains of unknown function (DUF). We thus examined several such multigene families for possible enrichment among candidates for individual trait groups. Our enrichment analyses included all seven DUF families represented by two or more genes in at least one trait group, as well as five large gene families containing multiple candidates associated with at least one trait group, namely the F-box proteins (568 paralogs in Arabidopsis), receptor-like protein kinases (307 paralogs), cytochrome P450 isoforms (256 paralogs), cysteine/histidine-rich C1 domain proteins (153 paralogs) and formins (FH2 proteins, 21 paralogs).

For nine out of these 12 gene families, we found in at least one category of GWAS candidates a reliable, significant at least 2x enrichment compared to whole genome abundance (Figure 8; Supplementary Table S9).These include receptor-like kinases (minor but significant enrichment in shape traits association), cytochrome P450 (enrichment in cell wall fluorescence-related traits, compare also Table 4), cysteine/histidine-rich C1 domain proteins (enrichment in shape and metal accumulation-related traits), formins, DUF674, DUF784 and DUF1985 proteins (enrichment in metal accumulation traits), as well as the DUF1261 and DUF3741 families that were found in association with trichome shape traits.

**Figure 8.**
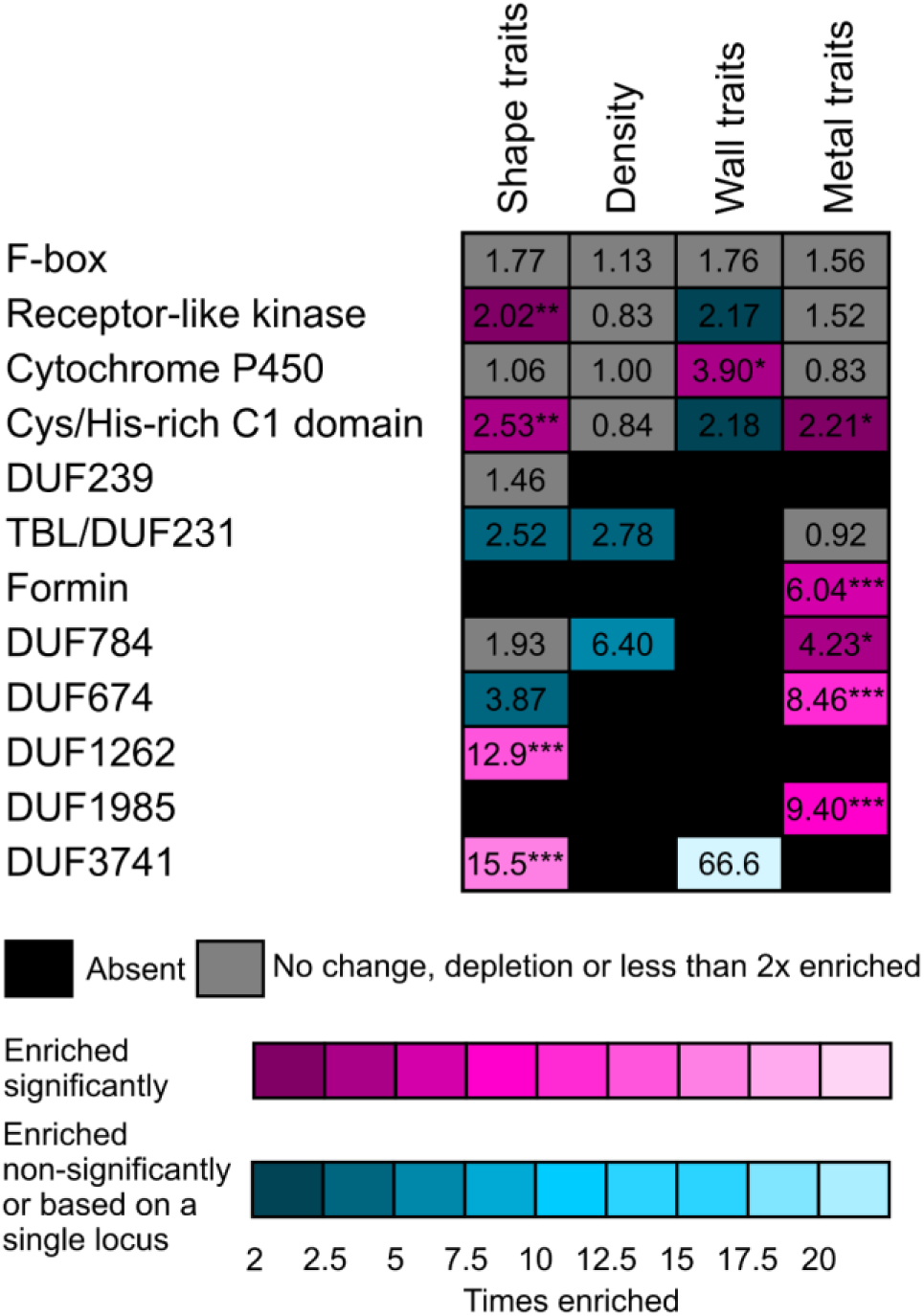
Summary of gene family enrichment analysis for selected gene families. The heatmap shows results of enrichment analysis for selected large gene families and all DUF (domain of unknown function) families that were represented by at least two candidates associated with at least one trait group. Numbers correspond to linear fold enrichment compared with the proportion of the family in question among all Arabidopsis loci (Araport11 genome annotation). Asterisks denote significant deviation from the proportion of the family in the whole genome (determined only for enrichment values of at least 2 using the Chi-square test with Benjamini-Hochberg correction for multiplicity; *** P<=0.001; ** 0.001<P<=0.01, * 0.01<P<=0.05).

Making use of available previously characterized loss of function mutants affecting all three formin genes associating with the stomatal metal deposition, i.e. *FH1*, *FH13* and *FH14*, we performed single-blinded evaluation of epidermal metal deposition in these mutants and corresponding wild type plants. The results suggest a tendency towards increased stomatal metal accumulation especially in the *fh1-1* T-DNA insertion mutant of the formin gene *FH1*, and possibly also in the CRISPR-generated loss of function allele of the same gene (Figure 9). Although none of the observed between-genotype quantitative differences was statistically significant, this observation is consistent with a possible contribution of formins to epidermal metal deposition that would deserve further attention. It also suggests that also the other observed gene family enrichments may be relevant.

**Figure 9.**
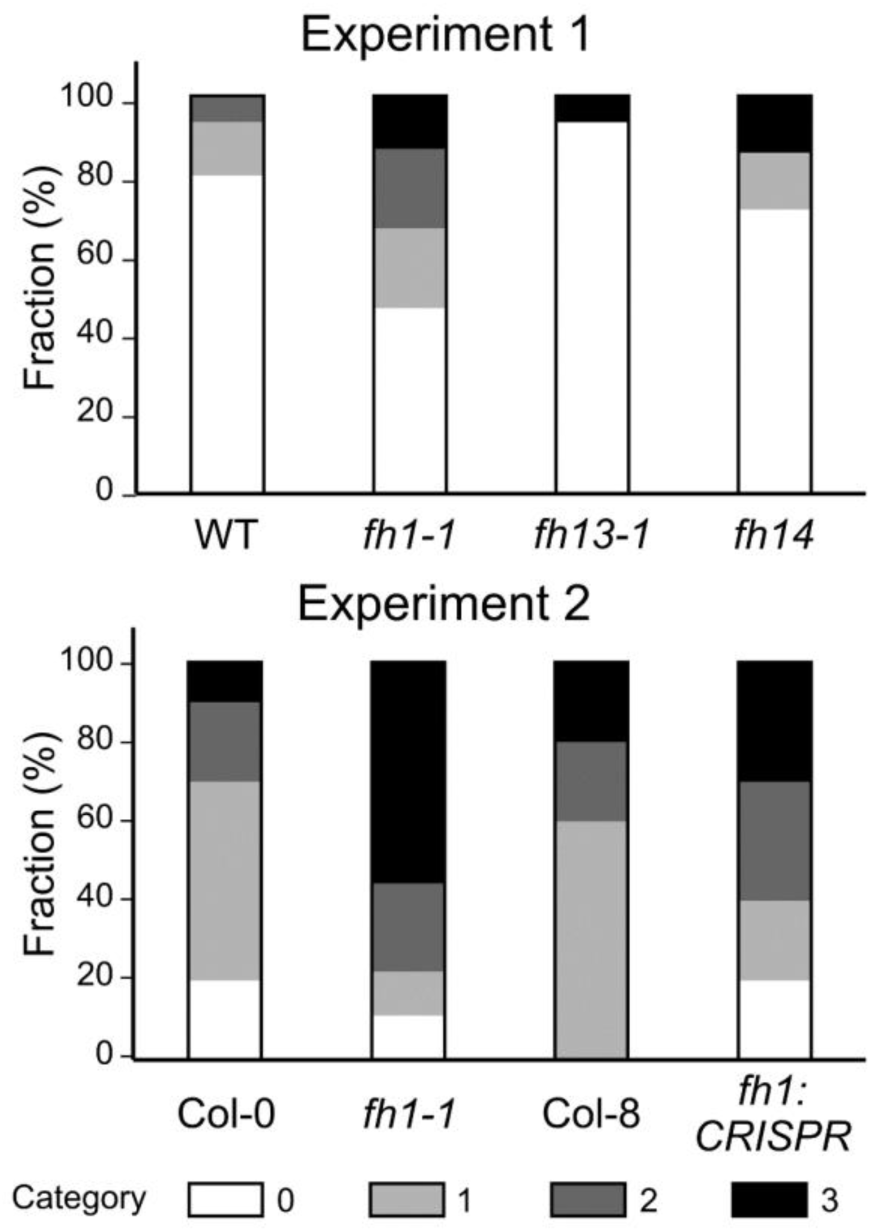
Stomatal metal deposition patterns in selected formin mutants and wild type plants. Data from two independent experiments, involving plants grown at different times under imperfect temperature and humidity control, are shown, each of them based on single-blinded evaluation of 9-15 plants per genotype for metal deposition patterns, categorized as in the original GWAS screen,. The wild type plants from experiment 1 were WT segregants from the Col-0 derived T-DNA mutant population that was the source of our *fh1-1*, *fh13-1* and *fh14* lines, while in the second experiment, *fh1-1* was compared to Col-0 and the *fh1:CRISPR* mutant to its progenitor genotype Col-8. The between-phenotype differences of interest were not statistically significant (Fisher-Freeman-Halton test P > 0.1).

### Associations among candidate genes indicate possible functional modules

To uncover functional relationships among genes associating with the studied traits, we scanned the candidate lists for mutual links recorded in the STRING database (Sklarczyk et al., 2023) that registers both experimentally documented and predicted interactions among genes, such as transcriptional co-regulation, clustering on the chromosomes, epistasis, or proteins, such as mutual binding.

Groups of associated genes were found for all four trait groups (Figure 10; Supplementary Figure S5, Supplementary Table S10), but only for trichome density and metal accumulation the numbers of interactions were significantly greater than predicted for a random gene selection (Supplementary Table S10). Inspection of the found clusters, however, reveals biologically meaningful associations also among candidates linked to the remaining trait groups. For example, among the shape-associated genes, cluster 15 consists of Exocyst complex subunits. For density and metal deposition, the clusters notably included mainly genes participating in nuclear functions such as chromatin organization, gene expression, RNA processing or nuclear transport.

**Figure 10.**
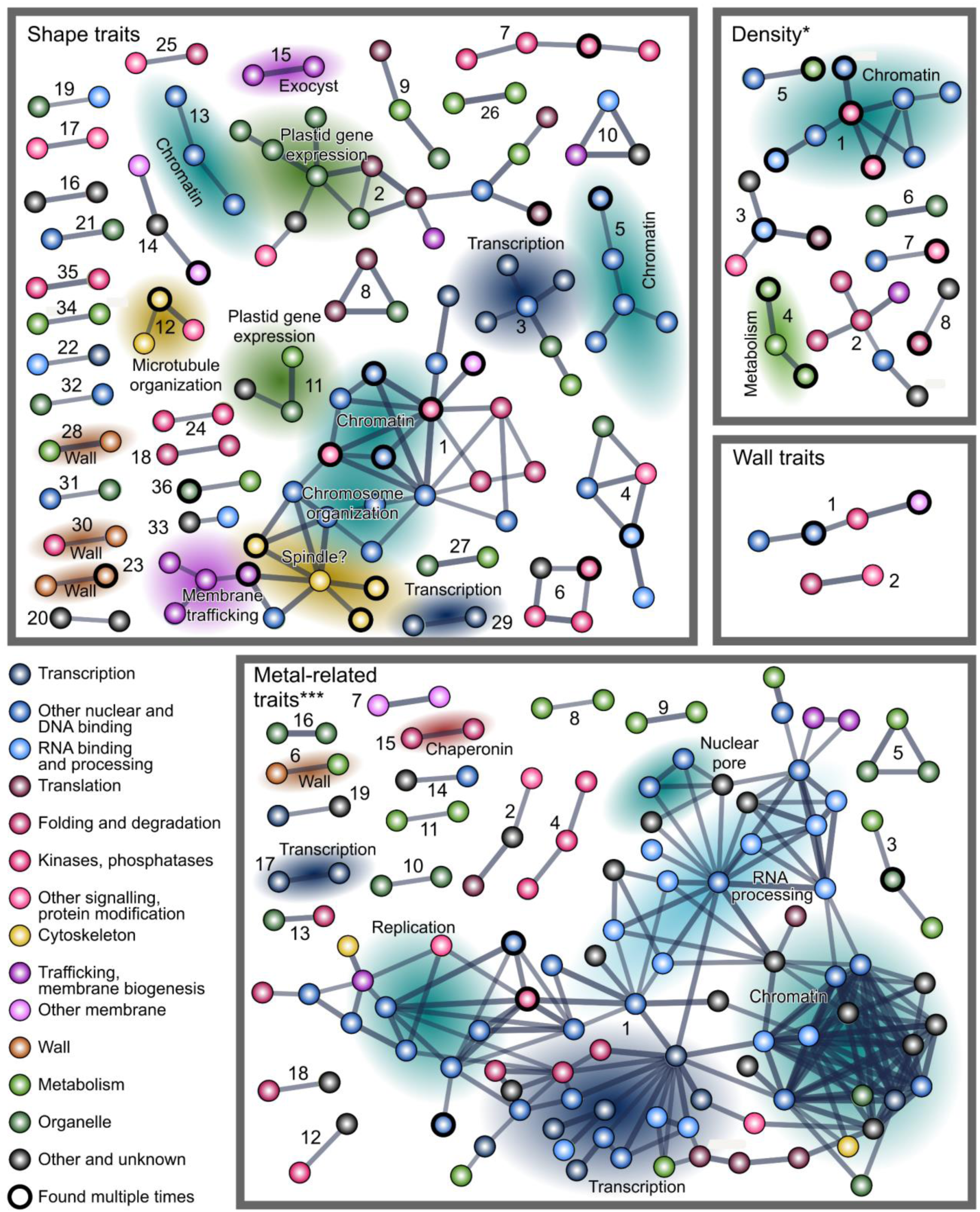
A schematic map of predicted interactions among candidate genes for individual trait groups. Thickness of graph edges reflects strength of evidence. Numbers correspond to the numbering of gene clusters in Supplementary Table S10. Asterisks denote significant enrichment of interactions compared to a random gene selection, as determined by the STRING algorithm (*** P<=0.001; * 0.01<P<=0.05).

Since we noticed that some genes are shared by clusters found for two or more trait groups, we next performed an analogous search for inter-gene associations among candidates associating with multiple phenotypic trait groups. With somewhat less stringent criteria than those used in the initial interaction searches, this group of genes also exhibited significantly more mutual associations than a random gene selection (Figure 11; Supplementary Table S10), with one large and two smaller clusters comprising mostly genes associated with shape traits, as well as a small cluster consisting of genes linked to trichome density. We believe that these findings may provide a starting point for exploration of novel regulatory pathways engaged in Arabidopsis epidermal differentiation.

**Figure 11.**
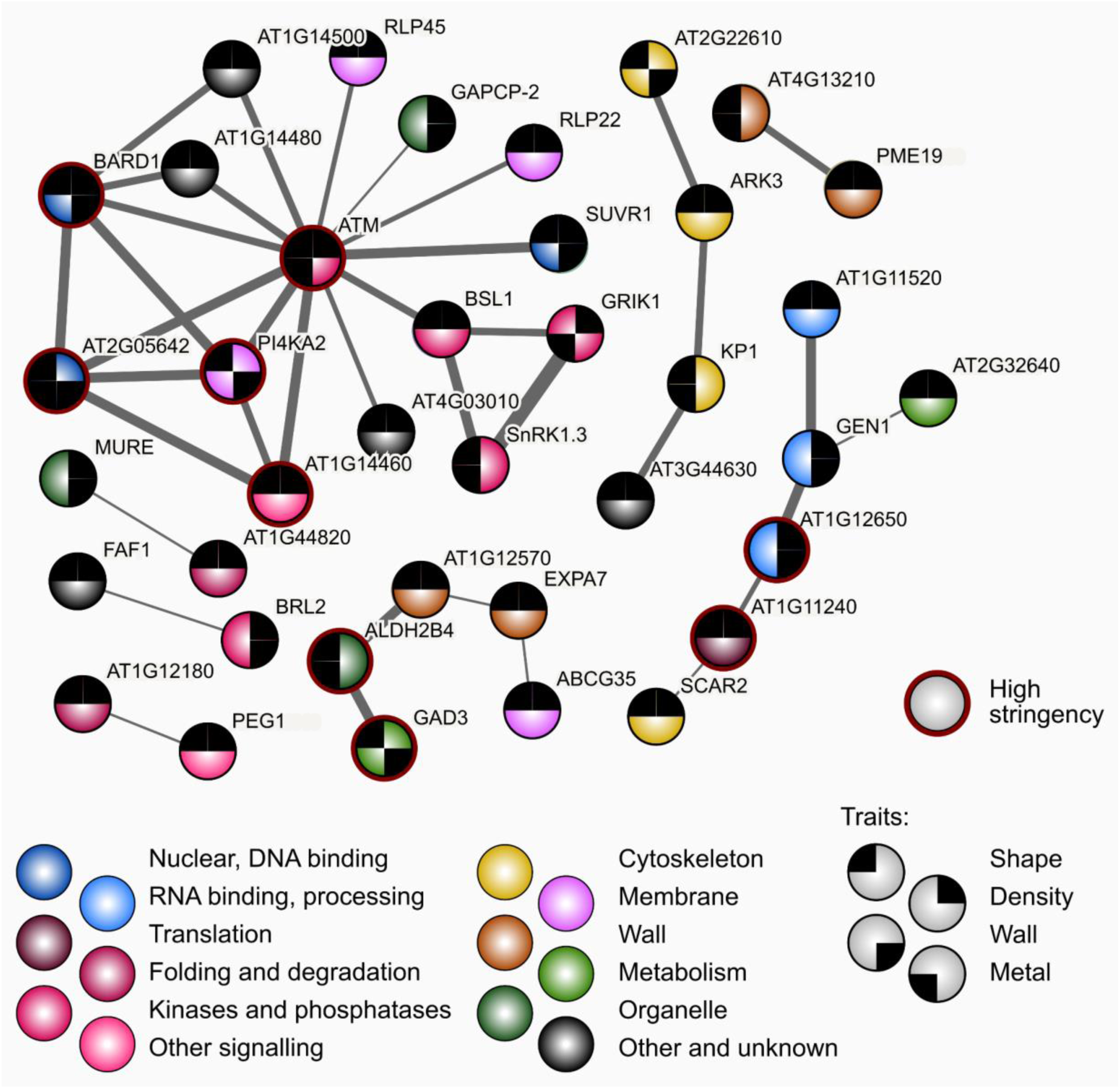
Predicted interactions among candidate genes linked to two or more trait groups. Black segments at graph nodes indicate the trait groups the gene was associated with. Note that the interactions were detected using lower stringency settings than those shown in Figure 9, but a part of the network was found also at high stringency settings (denoted by the red outline of participating nodes). Thickness of graph edges reflects strength of evidence.

## Discussion

We employed the GWAS approach to explore the genetic basis of *A. thaliana* phenotypic variability, focusing on epidermal traits including trichome development, secondary cell wall composition, production of autofluorescent substances, including those deposited in the cell wall, and metal accumulation. To our knowledge, no similar study has been conducted in Arabidopsis so far, except of a recent report (Arteaga et al., 2022) focusing on trichome patterning and trichome density (the latter also included in our study).

We studied 18 quantitative, semiquantitative or qualitative epidermal traits from 310 predominantly Scandinavian *A. thaliana* accessions from the 1001 Genomes collection (1001 Genomes Consortium, 2016). These traits can be divided into four groups (trichome shape, trichome density, cell wall composition and metal deposition-related). All quantitative parameters, as well as most semiquantitative categorical ones, exhibited continuous variability. As expected, traits reflecting trichome size, as well as those describing trichome shape complexity, were mutually positively correlated. Some of the analyzed traits exhibited also significant, albeit usually weak, correlation with selected climatic variables of the sites of origin of our plant accessions. Notably, epidermis autofluorescence was consistently weakly positively correlated with variables reflecting ambient temperature and amount of precipitation. Autofluorescence of plant tissues is largely due to phenolic compounds that might contribute to the defense of plants against microbial pathogens that thrive in warm, wet conditions (Donaldson, 2020; Bhattacharya et al., 2010). However, the only observed correlation reaching at least the moderate range (*sensu* Schober et al., 2018) was a negative one between maximum temperature of the warmest week and trichome stem length, where a biological interpretation was not immediately obvious.

Surprisingly, we noticed strong dithizone staining, presumably detecting zinc and other heavy metal ion accumulation (see, e.g., Frederickson, 2003; Srivastava et al., 2014; Peco et al., 2020), in stomatal guard cells of some accessions (but not in the common Col-0 line). A similar pattern was reported for Cd accumulation in cadmium-stressed Col-0 plants (Zeng et al., 2017). In metazoan cells and in yeast, zinc accumulation is associated with increased secretory activity (Yuan, 2011), and this also may be the case in the guard cells, which exhibit vigorous membrane turnover.

The outcome of GWAS studies always presents a compromise between specificity and sensitivity (see e.g. Brachi et al., 2011). We found statistically significant missense SNPs associated with phenotypic variability for 10 out of the 18 parameters examined, affecting over 5 % of all *A. thaliana* protein-coding genes. Moreover, none of our candidate genes corresponded to any of the 15 trichome density-associated loci found in the recent GWAS report (Arteaga et al., 2022), which was based on phenotyping 191 predominantly Iberian *A. thaliana* accessions. This raises concerns whether our screen was specific and sensitive enough to be informative. However, genotype collection size in both studies was obviously far below the saturation threshold. For the similarly sized human genome, GWAS saturation was achieved only with population size several orders larger (Yengo et al., 2022). Our list of trichome density-associated genes also did not contain any previously characterized trichome patterning genes from the list compiled by Arteaga et al., 2022), nor did we observe significant enrichment of genes encoding most abundant trichome transcripts (Jakoby et al., 2008) or proteins (Huebbers et al., 2022). In case of the metal accumulation traits, where most candidates reflected the guard cell staining, there was no enrichment of the most abundant guard cell transcripts (Leonhardt et al., 2004), or proteins (Zhao et al., 2008). It is not uncommon that candidate genes from GWAS studies are not enriched in, or even are depleted of, known pathway genes. This would be consistent with the assumption that mutations in large effects genes, which typically tend to be the known genes, might not be beneficial in natural environments (Ristova et al., 2018). Taken together, we believe that at least some of our candidates are biologically relevant, because the candidate genes set, both as a whole and for each trait group except metal accumulation, was significantly (or, in one case marginally significantly) depleted of transposon-borne loci, whose SNPs, unlike polymorphisms concerning transposon presence or position (see, e.g., Yan et al., 2022), are rather unlikely to have phenotypic effects. Thus, the sets of trichome shape, density and probably also wall trait-associated GWAS candidates are likely to contain biologically meaningful genes, while the relevance of candidates found on the basis of metal-related traits remains questionable.

We recovered five loci with well documented mutant phenotypes consistent with the associations identified in our study. First of them is the AT2G20190/CLASP gene coding for a microtubule-associated protein whose loss reduces trichome branch number (Pietra et al., 2013), In this gene, a SNP causing a conserved M1399L substitution in the C-terminal Armadillo-like repeat correlated with longer trichome stems. This substitution occurred more frequently in accessions from locations with higher Bio05 climate variable, reflecting the maximum environmental temperature at the site of the accession’s origin. CLASP stabilizes microtubules and consequently affects cell division and cell expansion (Ambrose et al., 2007; Pietra et al., 2013). The stability of Arabidopsis microtubule assemblies can be impaired by high temperatures, resulting in defective meiosis (De Storme and Geelen, 2020). Although a direct meiotic role of CLASP has not been documented, it is conceivable that the observed environmentally correlated allele distribution may reflect selection on the basis of a phenotype unrelated to epidermal development. This may be the case also for other loci with environmental bias in allele distribution, such as SYT1, which codes for a synaptotagmin involved in repair of membrane breaks caused by freezing (Yamazaki et al., 2008) and in defense against pathogens (Levy et al., 2015; Kim et al., 2016) whose abundance could be linked to environmental conditions.

The second candidate gene with a relevant mutant phenotype is AT2G38440/DIS3, encoding a subunit of the WAVE complex regulating Arp2/3 mediated actin nucleation. Its mutation causes a characteristic distorted trichome shape (Basu et al., 2005). Here, a conserved I1265V substitution in an area immediately preceding the WASP homology motif associated with reduced trichome circularity and solidity, as well as lower trichome density. The third gene, AT5G49830/EXO84B, codes for an exocyst complex subunit with a pleiotropic mutant phenotype that includes trichome deformation (Kulich et al., 2015; Fendrych et al., 2010). A non-conservative E123G substitution at a variable sequence position of this locus (see Cvrčková et al., 2012) associated with decreased trichome circularity and solidity. A non-conservative S93N substitution in a variable segment of another exocyst subunit, AT3G09520/EXO70H4, involved in metal deposition in the trichome cell wall (Kulich et al., 2015), associated with increased guard cell metal staining.

Last but not least, we found two conservative substitutions (S626A and D671E) associating with a decreae of trichome autofluorescence and also trichome density in AT3G28430/GFS9/TT9, which codes for a Golgi-associated peripheral membrane protein involved in membrane trafficking, vacuole development and flavonoid accumulation whose mutation results in altered autofluorescence (Ichino et al., 2014; Ichino et al., 2020). These mutations are frequently co-occurring with additional three substitutions, constituting a quintuple SNP minor allele, which appears to be present in multiple accessions of mountain origin across the globe. It is tempting to speculate that a variant with reduced cell wall autofluorescence may have altered flavonoid distribution among cell compartments, and perhaps enhanced vacuolar accumulation of anthocyanins that might bring a selective advantage under high altitude conditions characterized by increased UV stress, resulting in the fixation of this minor allele in mountain habitats. The presence of five identical SNPs in geographically disparate populations points towards this allele’s unique origin and subsequent elimination in most populations, rather than parallel independent mutations analogous to those found for example in naturally occurring “green revolution dwarf” Arabidopsis mutants (Barboza et al., 2013). The quintuple substitution allele of GFS9/TT9 would clearly deserve a follow-up experimental study.

Among the loci associated with trichome shape-related traits and also highly expressed in trichomes were found three genes participating in the biosynthesis of cutin and/or cuticular waxes. AT1G01600/CYP86A4 encodes a cytochrome P450 paralog, associates also with the trichome autofluorescence trait. Its expression is controlled by a transcription network that regulates also trichome branching (Camoirano et al., 2020; see also Berhin et al., 2022). Remarkably, the cytochrome P450 gene family is significantly over-represented among cell wall trait-associated candidates, also consistent with its role in cell wall biogenesis. The second trichome shape-related candidate with high trichome expression, AT1G12570, encodes a glucose-methanol-choline oxidoreductase related to the maize IPE1 gene participating in pollen exine development, which is co-regulated with CYP86A4 (Kannangara et al., 2007). Polymorphisms in both loci correlate also with variation in trichome density. Remarkably, trichome patterning is altered in some cuticle biogenesis mutants (Berhin et al., 2022), suggesting cuticle contribution to the mechanical cell coupling and tissue patterning (Reynoud et al., 2021). The third locus implicated in cuticle biogenesis is AT1G76690/OPR2, a 12-oxophytodienoic acid reductase co-regulated with the previous two loci (Suh et al., 2005), which only associated with variation of trichome stem length.

Relevant candidates were also found among guard cell-expressed genes. The loci associated with increased guard cell metal accumulation and highly expressed in guard cells include AT3G13772/TMN7, encoding a membrane protein whose heterologous expression causes yeast to accumulate copper (Hegelund et al., 2010), and AT5G39040/ALS1, coding an ABC transporter whose loss leads to increased aluminum sensitivity (Larsen et al., 2007).

Another gene implicated in stomatal metal accumulation, AT3G13460/ECT2, encodes a mRNA stability-controlling protein whose mutational loss affects trichome morphogenesis (Scutenaire et al 2018; Wei et al., 2018). Identification of this locus, which is intensively expressed in guard cells (Zhao et al., 2008) in the context of a guard cell-related phenotype indicates its broader role in epidermal development.

Eighty further candidates with likely roles in biologically relevant processes such as membrane trafficking, cytoskeletal organization or cell wall biogenesis were recovered. Among them are several members of the trichome birefringence-like (TBL) gene family whose founding member was discovered due to altered trichome cell wall optical properties (Bischoff et al., 2010), an additional exocyst subunit, multiple myosin- or kinesin-related proteins, several extensins and expansins, and about 20 cell wall modifying enzymes.

While these observations can be considered largely confirmatory, our candidate gene lists are significantly enriched in members from several multigene families not yet reported to participate in relevant aspects of epidermal development. The cysteine/histidine-rich C1 domain proteins, a group of nucleic acid binding proteins with a possible role in plant defense (Hwang et al., 2014) were enriched among trichome shape and metal accumulation-linked candidates, and receptor-like kinases (RLKs) associated with trichome shape traits, although no overlaps with a published list of trichome-expressed LRR-RLKs (Wu et al., 2016), representing one of the RLK family branches, were detected. Formins, evolutionarily conserved cytoskeletal organizers that also engage in endomembrane dynamics (see Cvrčková et al., 2024), surprisingly associated with stomatal metal accumulation, including the main housekeeping Class I formin FH1/AT3G25500 (Rosero et al., 2016; Oulehlová et al., 2019; Cifrová et al. 2020). Two candidate conservative amino acid substitutions were found in FH1 (L5F within the membrane insertion signal and D502E in a variable cytoplasmic segment). Observations in two loss of function *fh1* mutants suggest a trend towards increased metal accumulation in guard cells. Last but not least, members of five hitherto uncharacterized “domain of unknown function” (DUF) families associated with either trichome shape or metal accumulation traits, including the prolamin-related DUF784 (Zhang, 2009), suggesting their participation in relevant developmental pathways.

Although our Gene Ontology term enrichment analyses did not indicate links between the candidates and specific cellular functions, we observed a significant increase in mutual genetic, physical and functional interactions recorded in the STRING database (Sklarczyk et al., 2023) among candidates for the trichome density and metal deposition traits, as well as for candidates associated with multiple traits, compared to a random gene sample. Remarkably, the interacting gene clusters consisted predominantly of genes participating in nuclear functions. A prominent cluster of candidates associated with two or more trait groups comprising mainly signaling and regulatory proteins was centered on a homolog of the mammalian ATM kinase engaged in DNA damage response (Lee and Paull, 2021). In plants, ATM homologs participate in DNA repair and UV stress response (Shi and Liu, 2021), but also in reaction to exogenous DNA (Vega-Muñoz et al., 2023). Among genes associated with more than one trait category, we also found smaller clusters of shape-associated genes presumably engaged in cytoskeletal and cell wall-related functions, as well as an additional cluster comprising several nuclear proteins and containing genes linked to trichome density.

In summary, our screen identifies several candidate gene families and functional gene clusters that would deserve experimental study to verify their possible roles in epidermal development, possibly leading to uncovering new genetic players in cell morphogenesis and cell differentiation, but also in functions possibly relevant for environmental adaptation under natural conditions.

## Materials and methods

### Plant material

A set of 310 natural *Arabidopsis thaliana* accessions from the 1001 Genomes collection (1001 Genomes, 2016), has been included in our study (Supplementary Table S1). Predominantly, genotypes of North-European origin were chosen to reduce the effect of geographical differences in population structure and related parameters reducing GWAS efficacy (see Gloss et al., 2022). Climate variables data for the locations of origin vere retrieved in the form of standardized BioClim variables from the CliMond database (Kriticos et al., 2012).

Seeds were sterilized with chlorine gas, stratified in tubes at 4 °C for 3 days in the dark and sown into standard soil substrate pre-treated with Confidor to prevent blackfly infestation. Plants were grown in growth chambers under controlled long-day conditions (16 h light, 23 °C/8 h darkness, 18 °C; relative air humidity 60 %). Three seedlings per genotype were planted into individual pots at the first true leaf stage. Leaves were collected at the age of 5 weeks (at which point plants of some genotypes were beginning to bolt).

For mutant genotype verification for selected formin-encoding genes, homozygous plant lines carrying previously characterized loss of function mutants *fh1-1* (SALK-032981; Rosero et al., 2013), *fh13-1* (SALK_064291C; Kollárová et al., 2021), and *fh14-1* (SALK_058886; Li et al., 2010), available from NASC (RRID: SCR_004576), as well as *fh1:CRISPR* (generated in our laboratory, see Cifrová et al., 2020), with corresponding congenic wild type plants were used.

### Leaf sample processing

Three approximately fingernail-sized mature, non-senescent rosette leaves per plant (i.e. nine leaves per genotype) were harvested, resulting in three pooled samples, each containing one leaf from each individual. Samples were stored for at least two days before further processing or observation.

For trichome shape evaluation and for determination of callose content in trichomes, leaves were harvested into a tube containing 1 x PBS (phosphate-buffered saline) and 100 mM EGTA. Trichomes were isolated and stained for callose by aniline blue as described previously (Kulich et al., 2015). In genotypes where trichomes failed to detach from the leaves (further referred to as “shaveproof”), whole leaves were stained for callose using an analogous procedure (Kulich et al., 2018). This *in situ* staining was also used for additional documentation, such as the photos shown in Figure 1.

For visualization of autofluorescence and for determination of additional morphological parameters, leaves were pressed adaxial side up between two microscope slides and left to dry up to induce breakage of trichomes resulting in increase of autofluorescence (Kulich et al., 2018).

For visualization of metal content, leaves were collected into a tube containing 3 ml of acetone and stored for at least several days. On the day of observation, they were stained by addition of diphenylthiocarbazone (dithizone) reagent freshly prepared from 1.5 mg of dithizone, 1 ml of distilled water and 1-2 drops of glacial acetic acid (Seregin and Ivanov, 1997). After at least 1 h but not more than 6 h of staining, leaves were briefly washed in distilled water and mounted in water on microscopy slides for observation.

### Microscopy and imaging

Microscopic images were acquired using a Nikon Eclipse 90i fluorescence microscope equipped with PlanApo 4x/0.2 objective and Nikon DsFi 2 camera as described previously (Kulich et al., 2018). For autofluorescence and callose fluorescence detection, UV-B excitation was employed; in case of callose staining, an additional image was acquired in polarized light for trichome visualization. Leaves stained for metal detection were imaged using bright field settings.

If whole leaves were photographed, tilling images covering at least a half of the leaf lengthwise were automatically stitched from multiple frames, which were generated as maximum projections of 3 images with 50 μm of Z distance, using the Nikon Imaging Software (NIS Elements AR). Additional image processing was performed using the Fiji platform (Schindelin et al., 2012).

### Epidermal phenotypes determination

An overview of phenotypic trauts analyzed in our screen is provided in Table 1.

The first group of parameters was determined from photos of callose stained trichomes. At least 25 interactively selected clearly visible trichomes from 3 leaves of 3 plants (detached, or, in shaveproof genotypes, 8-9 in situ trichomes per leaf, selected across its diagonal) were evaluated. Trichome shape parameters (area, circularity, solidity, length and perimeter) were determined from binary images generated by thresholding polarized light photos using the Yen method as implemented in the Fiji software; the procedure was partially automated using recorded macros to increase throughput, with individual trichome selection being the only manual step. Trichome length was measured manually as the longest trichome dimension. Average parameter values from at least 25 trichomes were recorded for these parameters. Callose content was measured as the fraction of trichome area stained for callose, determined using built-in functions of Fiji from binary images generated by thresholding fluorescence and polarized light photos using the Yen method, again with the aid of recorded macros and manual selection of individual trichomes. The reported value for each accession is the sum of percentage values from 25 trichomes.

The second group of parameters was determined from fluorescence microscopy images of dried whole leaves, taken in the autofluorescence channel. Trichome stem length was measured using the NIS elements software with manual endpoint identification; average values from at least 25 clearly visible trichomes from 3 leaves, selected 7-9 per leaf across its diagonal, are reported. Additional parameters (trichome autofluorescence, overall leaf epidermis autofluorescence, autofluorescence color, trichome density and typical number of trichome branches) were visually categorized (see Table 1).

A third group of parameters reflects visually detectable patterns of metal accumulation, categorized as described in Table 1. This group includes the presence of a metal-enriched Ortmannian ring at the bottom part of the trichome stem (Kulich et al., 2015), presence of a visible metal staining gradient in the trichome stem, presence of metal staining at the trichome base, diffuse epidermal staining outside trichomes, and staining of stomatal guard cells.

The final categorical parameter, denoted as “shaveproof”, reflects the resistance of trichomes against mechanical detachment during staining for callose.

All categorical values were based on the typical appearance of the epidermis of three leaves. Glabrous leaves were attributed the value “0” for all categorical variables. In theexceptional cases where noticeable differences were seen among the three leaves of a given accession, the majority phenotype was recorded.

Phenotype data for a subset of accessions and traits have been independently rescreened by different team members for the purpose of broad sense heritability estimation (Supplementary Table S2); this also documented good inter-observer replicability of phenotype determination (Supplementary Figure S6).

### Phenotypic data processing and initial analyses

Primary phenotypic data were assembled into a structured spreadsheet and deposited in the public AraPheno database (Togninalli et al, 2020) as Study No. 126 (Bezvoda, 2023). Broad sense heritability (H2 = VG/VP), i.e. the proportion of phenotypic variation (VP) due to genetic variation (VG) estimated from the between- and within-line phenotypic variance, was calculated from trait values of all individuals in this dataset (Alonso-Díaz et al., 2021).

Statistical analysis of phenotypic data and phenotype/environmental parameter correlation analyses were carried out by the Pandas data analysis toolbox (Reback et al., 2021) and pair plots of scatterplots were created by the Seaborn visualization tool (Waskom, 2021) v. 0.11.2. Correlation strength is reported using criteria from Schober et al. (2018).

Principal component analysis (PCA) has been performed using the PAST software (Hammer et al., 2001) v. 4.11 using the correlation matrix method. All quantitative or semi-quantitative parameters were treated as ordinal variables, the qualitative “autofluorescence color” and “shaveproof” traits were handled as nominal.

### GWAS analyses and genomic data processing

Phenotype values for each trait were uploaded to the online GWAS application (Seren et al., 2012; Seren, 2024) and correlation analyses were run at 5 % FDR threshold with all possible combinations of input data transformations and statistical methods available with default settings except the minor allele count (MAC) value that has been lowered to 5. The 1001 Full sequence Dataset (TAIR v. 9) was used as the source of genotype information. After initial manual exploration of resulting Manhattan plots, the process of analyzing data was automated.

Raw result data files were downloaded manually and used for subsequent steps. All SNPs identified as significant by at least one method were considered. Additional information about each significant hit was acquired from the GWAS application by web scraping using the Python programming language v. 3.8 (Python Software Foundation, 2024) and driver for Google Chrome web browser (Google, 2024). Locus annotation was based on TAIR9.0 to maintain compatibility with the 1001 genomes dataset. All intergenic hits were discarded from subsequent steps of data processing. Hits were categorized according to the position of the SNP in corresponding gene (INTRON, NON_SYNONYMOUS_CODING, START_LOST, STOP_GAINED, STOP_LOST, SYNONYMOUS_CODING, SYNONYMOUS_STOP, UTR_3_PRIME, UTR_5_PRIME).

Annotation of loci that were subjected to further investigation was individually updated by manual searches of the Araport 11 genome annotation (Cheng et al., 2011) accessed via the BAR Thalemine portal (Pasha et al., 2020).

To determine the direction of phenotypic parameter changes for individual minor allele amino acid substitutions, lists of accessions carrying specific substitutions retrieved from the Ensembl resource (Yates et al., 2022) were used to identify subsets of our accessions carrying individual sequence variants. Average values of the phenotypic parameters of question were subsequently determined for each such variant.

### Geographic mapping of allele distribution

Lists of accessions carrying specific alleles of selected loci for the purpose of environmentally correlated allele distribution analyses and for map generation were generated from Ensembl data as described above. Geographic coordinate-based maps were created using Python Folium (Python Visualisation, 2024) or Google Maps tools.

### Enrichment and depletion analyses of GWAS candidate gene lists

To determine whether lists of candidate genes identified in the GWAS are enriched or depleted for specific gene groups, we used the Arabidopsis genome (Araport 11 version) and compared our data lists after manual annotation with the following gene lists.

As mobile element-derived genes, those with annotations containing the word “transposable” were considered.

As genes with high transcript levels in trichomes, we considered the 164 genes listed as encoding 5 % most abundant transcripts in the mature trichome transcriptome (Jakoby et al., 2008).

As genes with guard cell-specific transcription patterns, we considered the list of loci specifically upregulated in the guard cells compared to the mesophyll (Leonhardt et al., 2004).

As genes encoding trichome-abundant proteins, we selected genes from the recently published Arabidopsis trichome proteome study (Huebbers et al., 2022) using the following criteria. A protein had to be significantly enriched in at least two of the four reported proteomic experiments, at least in one case by a factor of two and more, while it was not depleted in any experiment. These criteria produced a list of 455 loci.

For genes encoding guard cell-abundant proteins, we considered the list from a published guard cell proteome study (Zhao et al., 2008).

For gene family enrichment, we included all gene family members as identified by keyword search of candidate gene annotations. Family member counts were based either on literature, as in the case of F-box proteins (Kuroda et al., 2002), on TAIR (Reiser et al., 2024) Gene Families annotations (as in the case of receptor-like protein kinases and FH2 proteins), or estimated by keyword searches of the Araport 11 annotations (for all DUFs).

Genes from each list were identified among the GWAS candidates and significance of any observed enrichment or depletion was evaluated using pairwise Chi-square test (Stagroom, 2024) with Benjamini-Hochberg correction for multiplicity performed using an online calculator (Radua et al., 2024).

### Verification of candidates

Loss-of-function mutants in candidate genes were grown alongside corresponding wild type plants and leaf samples were harvested, processed and imaged as described above for the main ecotype screen. To minimize effects of observer’s expectation bias, a single blinded experimental design was employed, with one member of the team performing the imaging and others quantitatively evaluating microphotographs with coded labels without knowledge of the plant’s genotype. After decoding the identity of the samples, significance of between-genotype differences in the relevant categorical parameters was estimated by the Chi square test (or Fisher’s exact test in cases where some categories exhibited zero counts) using online calculators (Stagroom, 2024; Vasavada, 2016).

### Mapping of protein-protein interactions among candidature gene products

To gain initial insight into the interactions among candidate genes, the full network STRING protein-protein interaction database v. 11.5 (Szklarczyk et al, 2023) was searched with non-redundant lists of genes associated with the indicated trait groups as queries, using high confidence (score = 0.7) and high stringency (FDR = 1 %) settings. Statistical significance of interaction enrichment was obtained during this search. Resulting interaction networks were exported both as tables (further processed in Excel to generate cluster and node lists that were subsequently annotated as described above) and as vector images that were manually edited (to remove unlinked modes, tidy up the layout), annotated and colored.

Mapping of interactions among the subset of candidates associated with multiple trait groups was performed analogously except that the confidence threshold was lowered to medium (score = 0.4), i.e. to the default settings of the STRING database search tool.

## Acknowledgements

We thank Christian Göschl and Samantha Krasnodebski (Gregor Mendel Institute) for help with experiment setup and cultivating plants for the GWAS screen, Marta Čadyová for technical support, and an anonymous reviewer for very inspiring suggestions of additional data analyses.

We acknowledge continuous support from the Charles University Progres Q43 and COOPERATIO programs. The initial stages of this work have been supported by the CSF/GACR/FWF project GF16-34887L (V.Ž., W.B., Y.M.L-R., F.C.), the formin mutant verification experiments by the CSF/GACR 22-33471S grant (E.K. and F.C.), and the finalization of this report by the project TowArds Next GENeration Crops, reg. no. CZ.02.01.01/00/22_008/0004581 of the ERDF Programme Johannes Amos Comenius.

## Author Contributions

R.B. analysed the primary phenotypic data, co-ordinated data management during the project, and drafted most tables, data-based illustrations, and parts of the manuscript, Y.M.L-R, Z.K., E.K., I.K. and F.C. collected phenotypic data during the initial screen, Z.K. and Y.M.L.-R drafted photograph-based figures and parts of the manuscript, F.C. and R.B. rescreened primary data for replicability and heritability estimation, E.K. and F.C. performed phenotypic characterization of formin mutants, W.B. and V.Ž. conceived the project, developed the study design and participated in data analyses and interpretation, F.C. participated in developing the study design, performed PCA, gene family enrichment and interaction network analyses, participated in other data analyses and results interpretation, drafted the manuscript and performed final editing of figures, tables and supplementary materials.

## Supplementary tables (MS Excel *.xlsx file)

**Table S1. List of *A. thaliana* accessions included in this study.** Accession numbers refer to the 1001 Genomes collection (1001 Genomes Consortium, 2016). Geographical coordinates, altitude and selected climate (BioClim) variables of the site of origin are listed for each accession (with legends in a separate spreadsheet).

**Table S2. Individual plant data for a subset of traits and accessions.** These data were used for replicability and heritability estimation.

**Table S3. Broad sense heritability of selected traits.** The values shown were calculated from data obtained in an independent rescreening of a subset of parameters (see Table S2 and Materials and Methods).

**Table S4. Mutual correlation of epidermal traits and climatic parameters.** Separate spreadsheets are provided for Spearman’s correlation coefficients (Rho) and FDR-corrected P-values for these correlations. Values are color-coded as in Figure 4 and Supplementary Figure S1.

Table S5. List of all significant SNPs detected in this study.

**Table S6**. **Annotated list of all loci containing ORF-changing SNPs significantly associated with individual traits.** Separate spreadsheets are provided for individual parameters.

**Table S7. Annotated lists of all loci with ORF-changing SNPs significantly associated with multiple traits and trait groups.** Separate spreadsheets are provided for individual trait combinations.

**Table S8. Candidate loci whose annotation suggests a possible role in known processes affecting the relevant traits.** Processes involved include, e.g., cell expansion, cell wall biogenesis, tissue patterning or metal accumulation. Only loci not identified by gene expression patterns (see Table 4 and Table 5) are listed.

**Table S9**. **Numbers and lists of candidates from selected gene families.** Known large gene families and all DUF (domain of unknown function) families that were represented by at least two loci associated with at least one of the studied trait groups are listed, together with results of enrichment analysis including statistical evaluation.

**Table S10. An annotated list of mutually associated gene clusters.** Clusters and genes participating in them are listed together with results of interaction enrichment analysis including statistical evaluation.

## Supplementary figures (Portable document format *.pdf file)

**Figure S1. Mutual correlation of all epidermal phenotype parameters analyzed.** Distribution of values for each parameter is shown at the diagonal of the matrix, with accession counts on the Y axis. Spearman’s correlation coefficients (Rho), shown in the upper right triangle for all parameter pairs, are color-coded: gray for negligible correlation (−0.1 < Rho < 0.1), magenta for positive, cyan for negative correlation, with intensity of coloring reflecting correlation strength, while P-values for these correlations, shown in the lower left triangle, are color-coded from green (highly significant, 0 < P ≤ 0.01 with darker color corresponding to lower P) through yellow (significant, 0.01 < P ≤ 0.05) to orange (P > 0.05, not significant).

**Figure S2. Distribution of SNP types identified as significantly associated with individual parameters**. Only traits with detected polymorphisms in more than 5 loci are shown.

**Figure S3. Geographic allele distribution for selected genes with significantly environmentally correlated polymorphism.** Sites of origin of accessions harboring the reference allele are shown in blue, those with the minor allele in red (for allele description see Table 4, Table 6 and Figure 7A).

**Figure S4. Distribution of all standard climatic variable values for the reference and quintuple substitution minor allele of GFS9/TT9.** Data for whole Ensembl accessions set are shown. Accessions where some of the SNPs characteristic for the quintuple allele are missing due to sequencing gaps (“probable quintuple”) are included in the quintuple mutant category. FDR-corrected P-values for the observed differences are color-coded green for highly significant, 0 < P ≤ 0.01 (with darker color corresponding to lower P), yellow for significant (0.01 < P ≤ 0.05), and orange for not significant (P > 0.05). Red lines indicate median values.

**Figure S5. A detailed map of predicted associations among candidate genes for individual trait groups.** Thickness of graph edges reflects strength of evidence. Numbers correspond to the numbering of gene clusters in Supplementary Table S9.

**Figure S6. Inter-observer replicability of selected parameters estimation**. For continuous variables, results from two independent screens of the same images are plotted against each other (X-axis values correspond to averages of three plants). For categorical ordinate variables, the replicability values correspond to the percentage of accessions where results for at least two out the three plants differed by no more than 1 between two independent visual screens of the same images.

