## Supplemental Figures for "A genome-wide association screen for genes affecting leaf trichome development and epidermal metal accumulation in Arabidopsis"

**Figure S1. Mutual correlation of all epidermal phenotype parameters analyzed.**

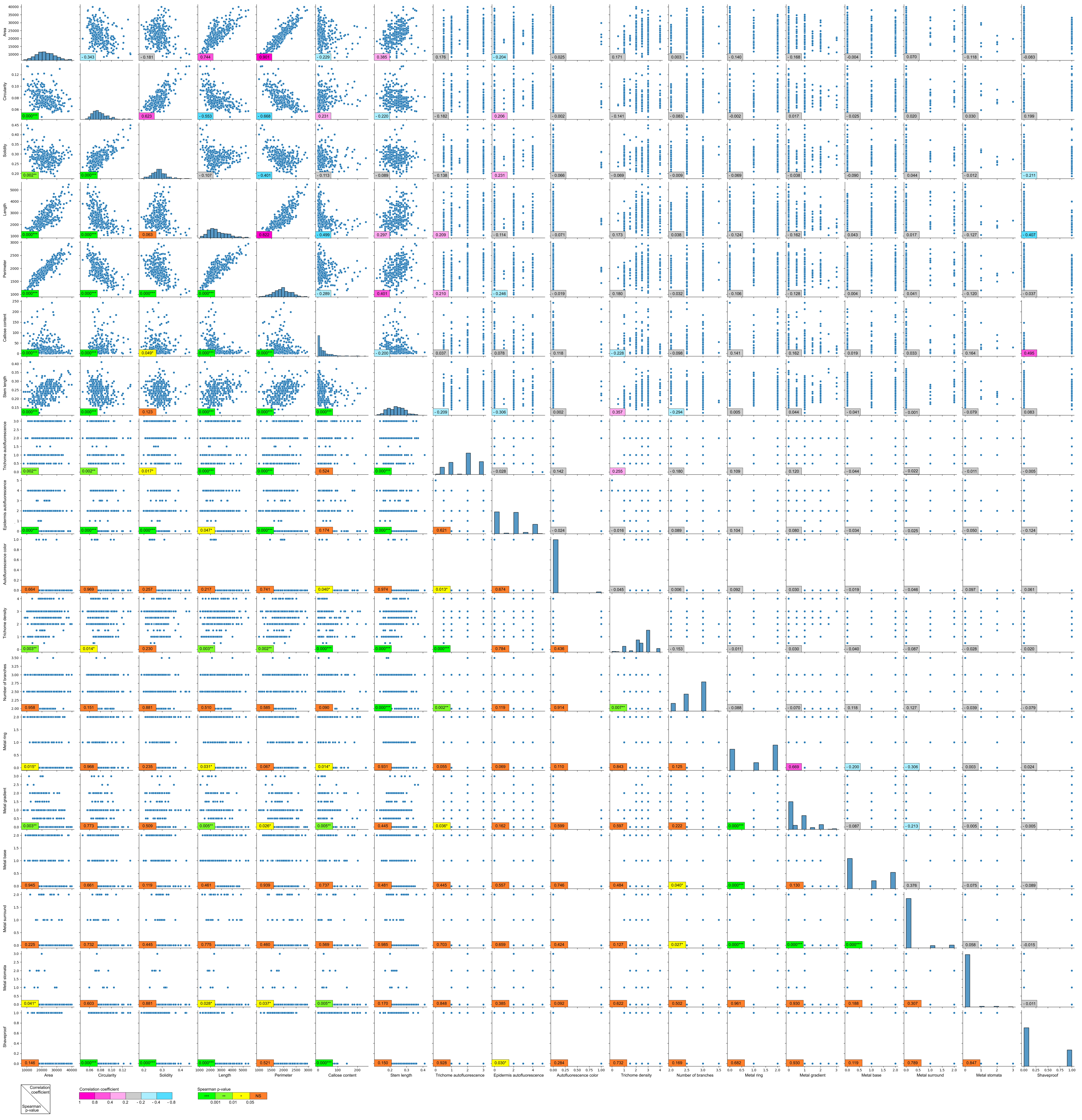

**Figure S2. Distribution of SNP types identified as significantly associated with individual parameters**.

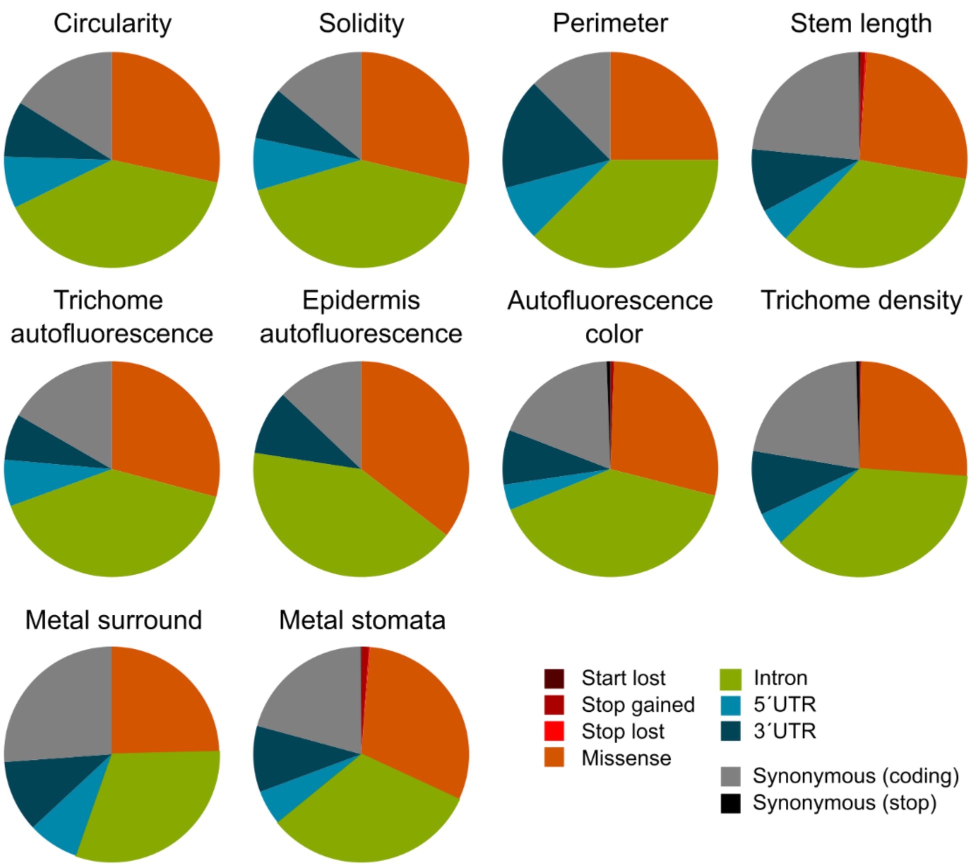

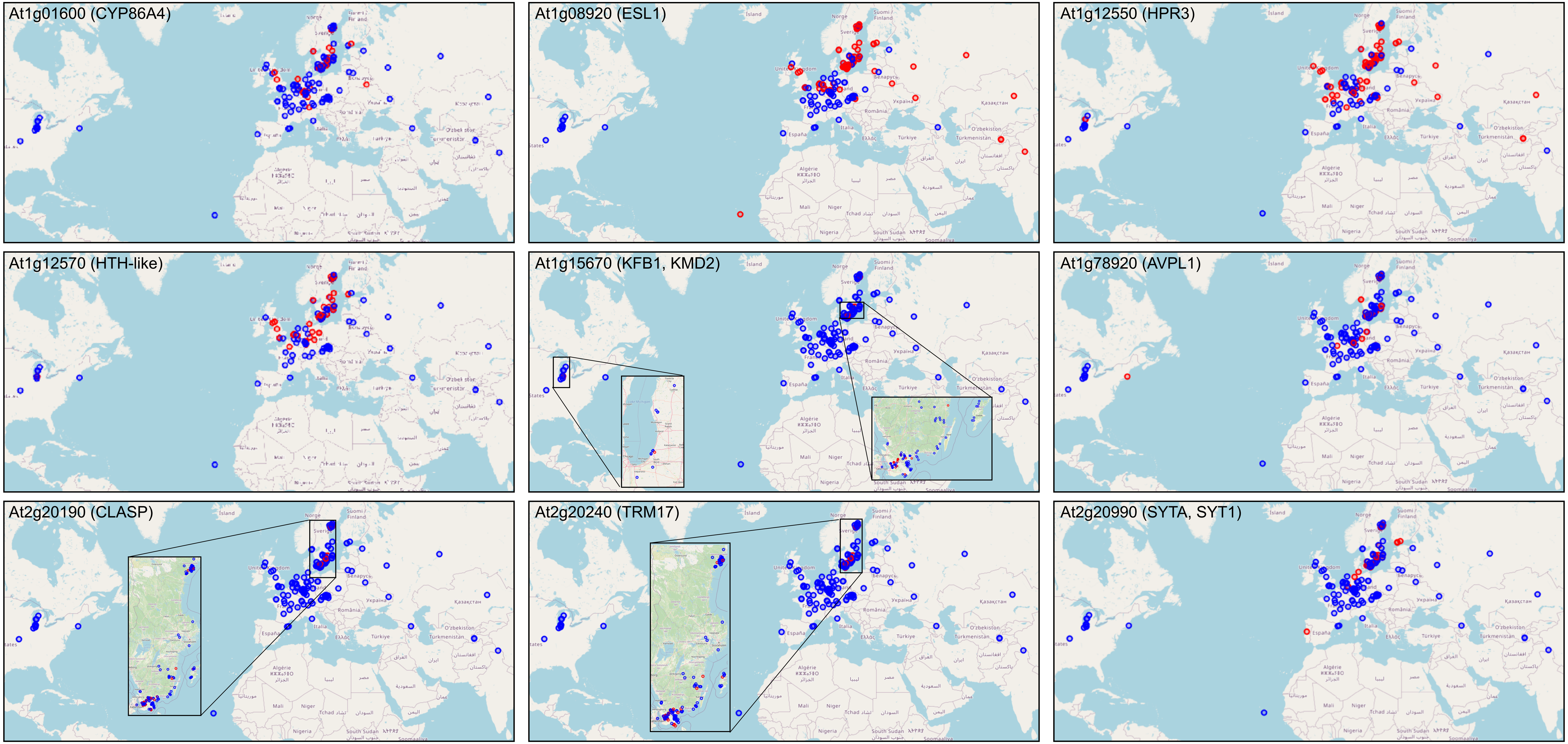


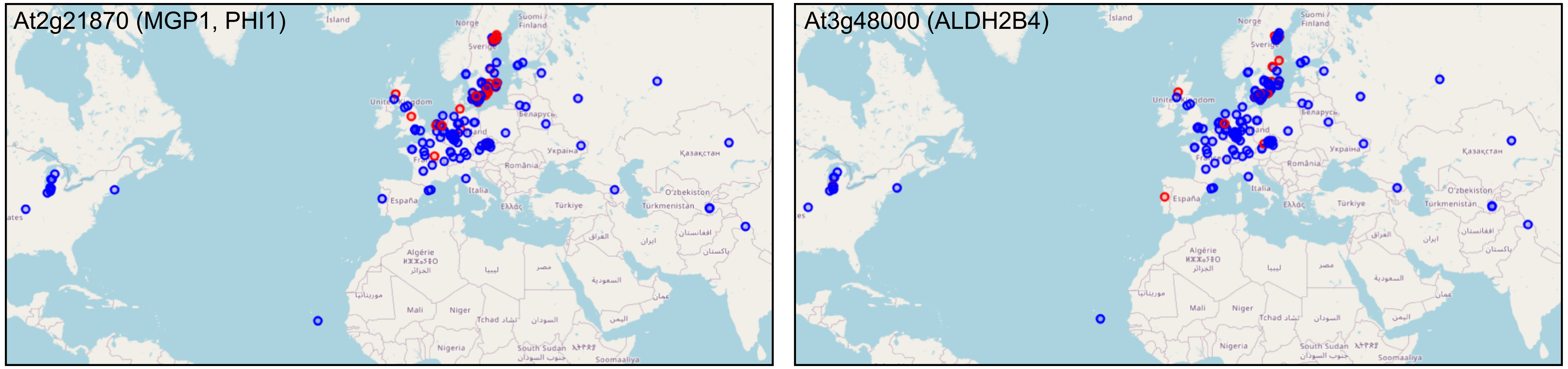


**Figure S3. Geographic allele distribution for selected genes with significantly environmentally correlated polymorphism.**

**Figure S4. Distribution of all standard climatic variable values for the reference and quintuple substitution minor allele of GFS9/TT9.**

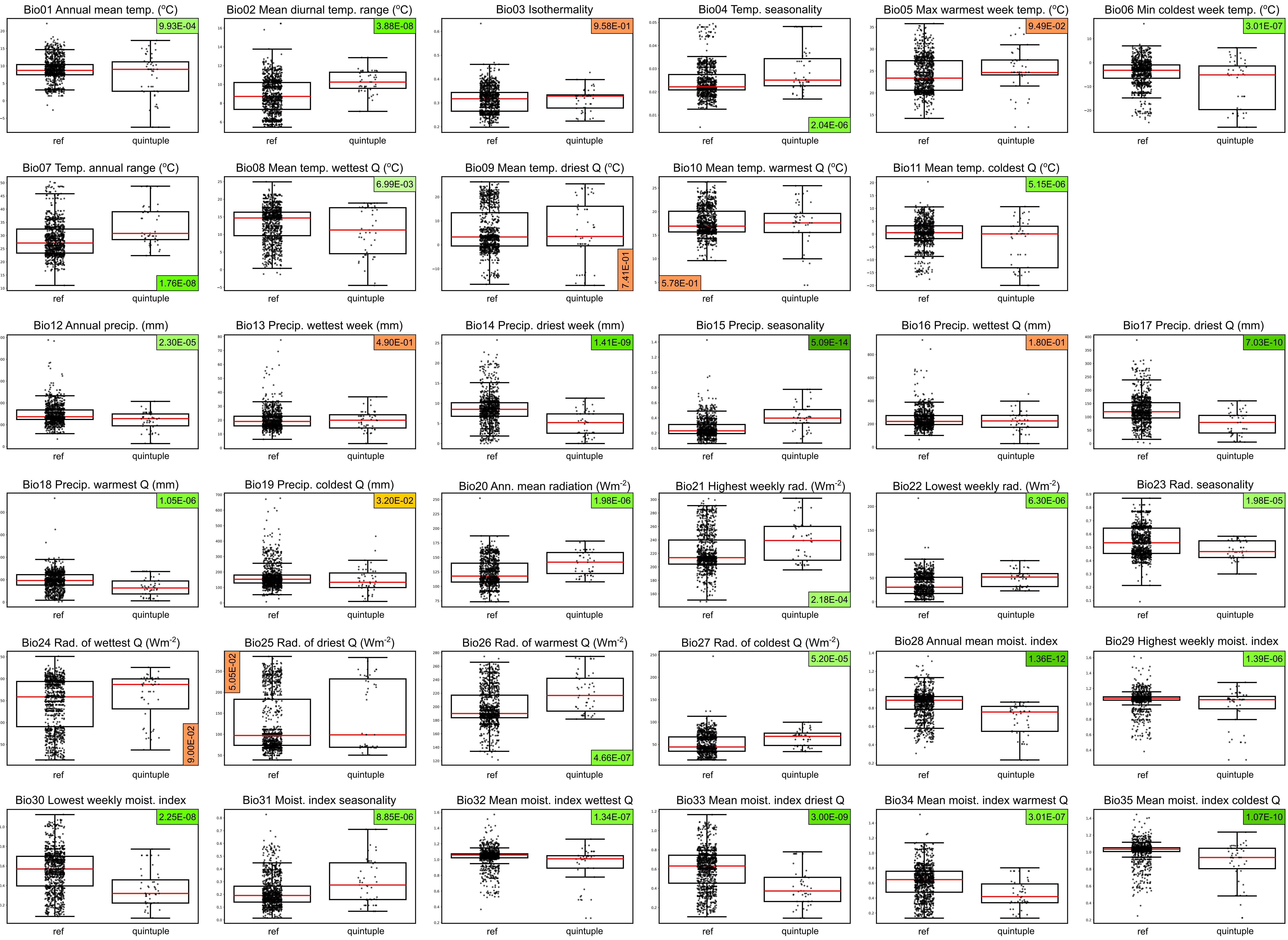

**Figure S5. A detailed map of predicted associations among candidate genes for individual trait groups.**

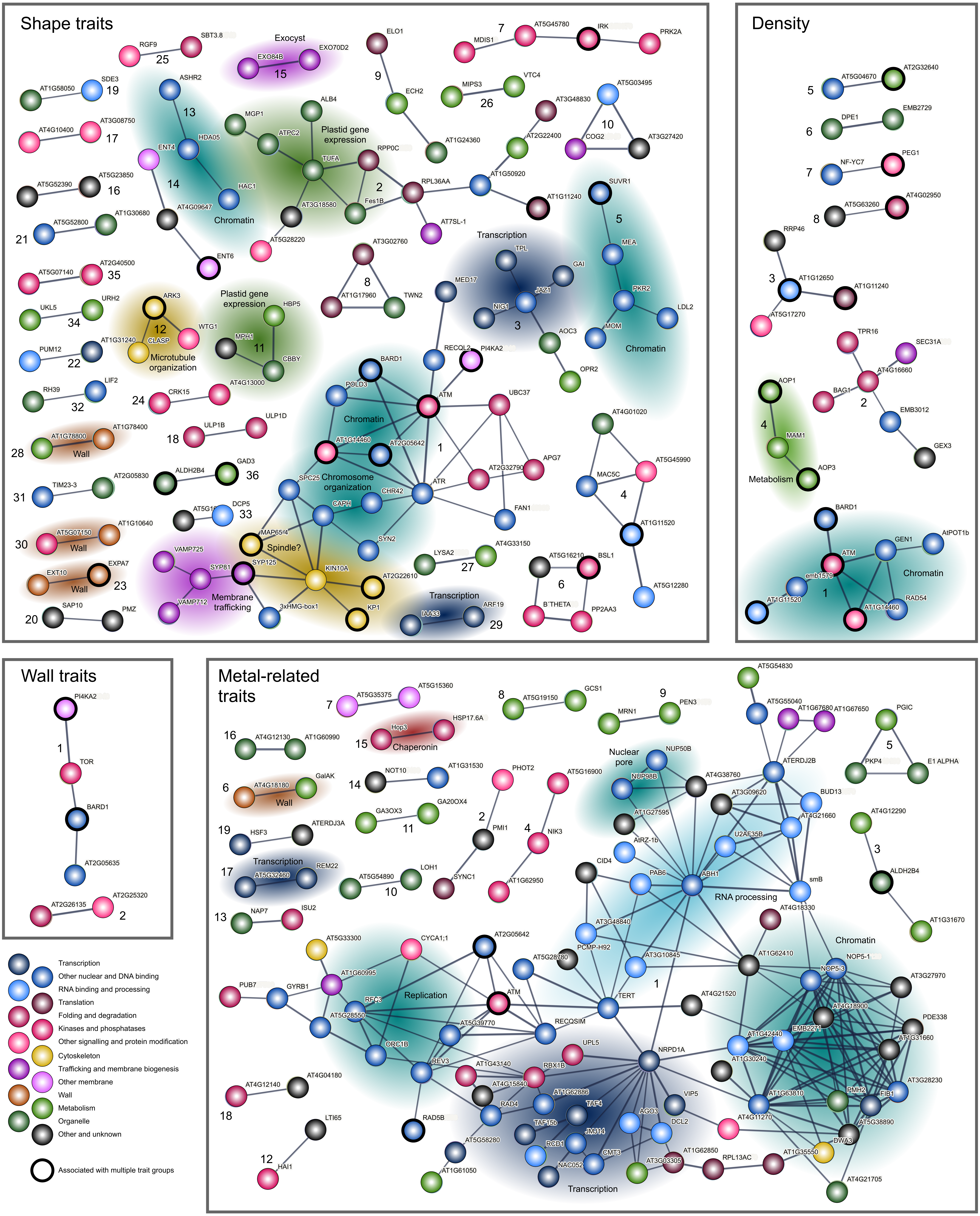

**Figure S6. Inter-observer replicability of selected parameters estimation.**

Area

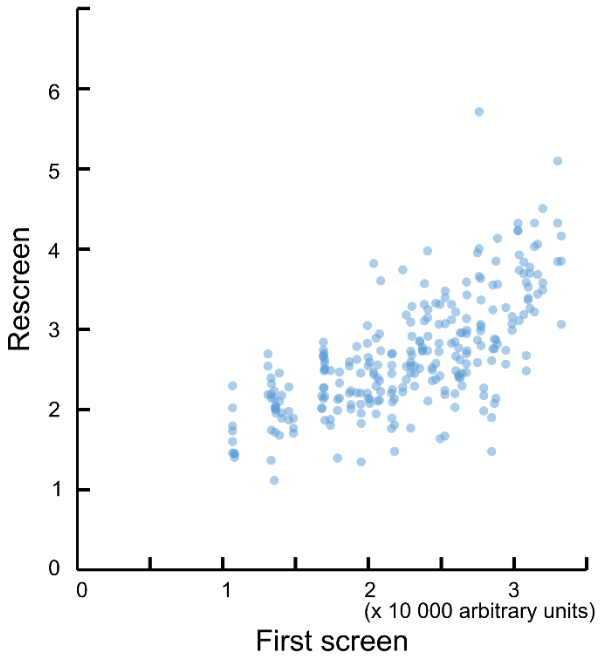


Callose content

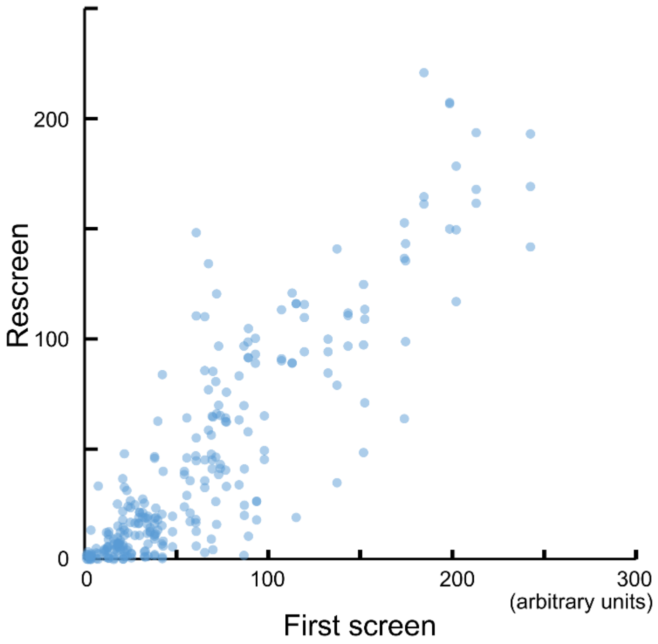


Stem length

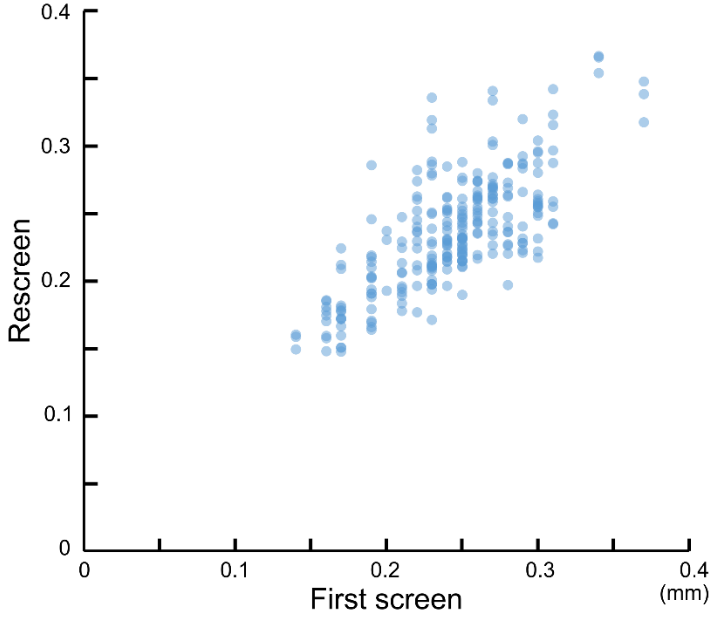


Categorical variables

| Parameter | Accessions rescreened | Replicability (%) |
|---|---|---|
| Trichome autofluorescence | 89 | 99 |
| Epidermis autofluorescence | 89 | 99 |
| Density | 90 | 98 |
| Number of branches | 89 | 97 |
| Metal ring | 88 | 100 |
| Metal gradient | 88 | 99 |
| Metal base | 88 | 98 |
| Metal surround | 88 | 95 |
| Metal stomata | 88 | 100 |
